# A Patient-Specific 3D Printed Carotid Artery Model Integrating Vascular Structure, Flow and Endothelium Responses

**DOI:** 10.1101/2025.06.29.661865

**Authors:** Jorge A. Catano, Louis Jun Ye Ong, Mingyang Yuan, Yunkun Qu, Jessica Benitez, Prasad KDV Yarlagadda, Yi-Chin Toh, Zhiyong Li

**Author notes:** Co-corresponding authors: Yi-Chin Toh, Address: 60 Musk Ave, Kelvin Grove, QLD 4059, Australia, Zhiyong Li, Address: 2 George St, Brisbane City QLD 4000, Australia. (J.A.C); (L.O); (M.Y); (Y.Q); (J.B); (P.Y); (Y.C.T); (Z.Y.L).

## Abstract

The progression of atherosclerosis is driven by the interplay between vascular anatomy, hemodynamic forces, and endothelial responses. However, existing in vitro vascular models have yet to integrate all these elements into a cohesive, patient-specific system. Here, we present the first instance of direct 3D printing of a miniaturized, patient-specific carotid artery model that recapitulates anatomical-dependent hemodynamic changes and vascular cell remodeling. Phase-contrast MRI scans from a healthy donor were used to generate miniaturized 3D carotid artery models, which were analyzed via computational fluid dynamics (CFD) and particle imaging velocimetry (PIV) to validate the preservation of physiological hemodynamic properties. Using digital light processing (DLP) 3D printing, we fabricated the miniaturized carotid artery vessel using GelMA, containing embedded human aortic smooth muscle cells and an endothelialized lumen. Perfusion culture replicated physiological arterial shear stress of up to 10 dynes cm^-^², resulting in differential endothelial cell alignment and inflammatory monocyte adhesion corresponding to laminar and turbulent flow regions within the carotid artery. This model serves as a powerful platform to investigate how anatomical variation influences susceptibility to atherosclerosis through its impact on local flow dynamics.

## 1. Introduction

Cardiovascular diseases (CVD) are the leading cause of mortality worldwide, often becoming clinically apparent during advanced stages of atherosclerosis, which leads to arterial blockages and thrombotic complications.[1] The development and localization of atherosclerotic lesions arise from a complex interplay between vascular anatomical structures, hemodynamic forces, and endothelial responses,[2] such as lipid accumulation, inflammation, and immune cell recruitment. Notably, bifurcation and branching regions within blood vessels are known to create recirculation zones, which reduce flow velocity and alter shear stress distribution along the vessel wall.[3, 4] These hemodynamic changes significantly affect endothelial cell function, promoting conditions favorable to atherosclerosis.[5–7] Furthermore, inter-patient variations in vascular structure and flow dynamics are closely associated with the propensity for adverse CVD manifestations, including ischemia and plaque rupture.[8] Developing comprehensive models that fully replicate these complex interactions will provide critical insights into the roles of anatomical and hemodynamic variables in driving atherosclerosis within specific vascular geometries.

To date, in vitro vascular models that can incorporate human-specific parameters are often limited to examining the crosstalk between pairs of factors, such as vessel structure and hemodynamic flows [9, 10] or hemodynamic flow and endothelial cell responses.[11, 12] Biofabrication techniques such as microfabrication, hydrogel molding,[13–15] electrospinning, and coaxial extrusion,[16] have successfully demonstrated the impacts of fluid shear stress on endothelial cell morphology and function. These models have also shown how endothelial cells respond to atherogenic environmental factors using patient-specific cells and exposure to proinflammatory cytokines and lipids.[17] However, the biomechanical environments in these in vitro models are often simplified, typically focusing on single parameters, such as shear stress or radial force, in geometrically simplified rectangular or cylindrical vessels.[18–22] On the other hand, elastomeric vascular phantoms have successfully replicated some patient-specific vessel geometries and mimicked the resultant complex hemodynamic conditions.[23] However, vascular phantoms are limited in replicating vascular tissue architectures needed for cellular studies.[24]

3D bioprinting has emerged as an attractive fabrication approach to construct vascular models, offering high precision in replicating 3D structures with excellent cell viability.[25, 26] Significant efforts have been dedicated to extrusion-based bioprinting, although this printing modality faces limitations in generating perfusable vessel structures since bioinks for extrusion printing often lack mechanical integrity.[27] To overcome this, sacrificial bioinks, such as pluronic acid, are bioprinted within cell-laden hydrogel blocks and later evacuated to create channels for medium perfusion.[28] The emergence of light-based 3D bioprinting techniques, such as digital light processing (DLP), has recently made it possible to fabricate perfusable, cell-laden vessel constructs with intricate 3D geometries. For example, Ching et al. demonstrated the use of a DLP-printed porous poly(ethylene glycol) diacrylate (PEGDA) mold to release a crosslinking agent for casting hydrogel vessel constructs, which successfully supported the co-culture of endothelial and smooth muscle cells.[29] Advances in the biocompatibility of 3D printing photo-resins, such as gelatine methacrylate (GelMA) photoinks, now enable the direct printing of cells into patient-specific vascular geometries.[30–32] Despite these advancements, current models have primarily focused on demonstrating the feasibility of supporting functional cell growth and endothelial layer formation under perfusion culture conditions. To date, no studies have shown that these 3D-printed patient-specific vessel models can replicate the complex interplay between vessel geometry, flow dynamics, and endothelial cell behavior.

In this study, we present the direct 3D printing of a patient-specific carotid artery model to investigate the interplay between anatomically dependent hemodynamic forces and vascular cell remodeling. Phase-contrast MRI scans from a healthy donor were used to generate a miniaturized 3D carotid artery model, which preserved physiological hemodynamic properties. This computational model was translated into a perfusable GelMA-based carotid artery model using DLP 3D printing, which contained embedded human aortic smooth muscle cells and an endothelialized lumen. Using this model, we show that we could study how patient-specific hemodynamic-induced vascular remodeling processes can contribute to atherosclerosis development and progression.

## 2. Results

### 2.1. Preservation of physiological hemodynamic flow in a miniaturized carotid artery model

To develop a reproducible and cost-effective vascular model that is biologically responsive for experimental use, we first assessed the feasibility of preserving physiological hemodynamic flow profiles in a patient-specific carotid artery model that has been miniaturized. Our approach involves using similitude analysis to predict the operating flow velocity required to maintain the physiological wall shear stress in a predetermined scaled-down model of a human carotid artery. Computational fluid dynamic (CFD) simulation was then performed to fine-tune the operating flow velocity in the miniaturized vessel model such that the shear stress distributions were closely aligned with those in full-scale simulations (Figure 1A).

**Figure 1.**
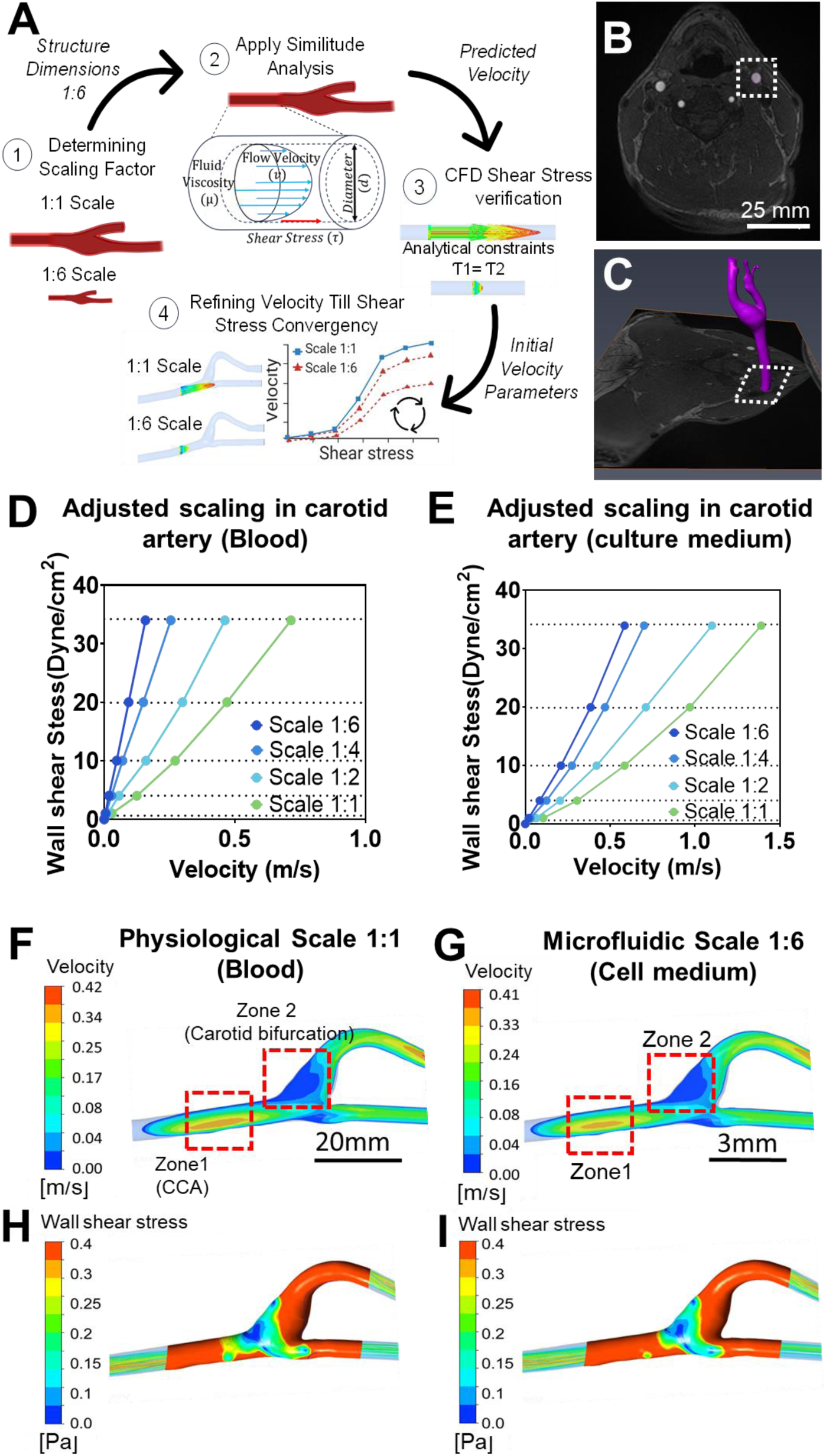
Geometrical and hemodynamic scaling of a carotid artery using analytical and computational analysis. (A) Systematic workflow to computationally replicate hemodynamic behavior within a scaled-down version of the carotid artery. (B) Time-of-Flight (TOF) cross-sectional image highlighting the carotid lumen. (C) Reconstructed carotid artery geometry derived from segmented cross-sectional image stacks. (D-E) Equivalent wall shear stress (WSS) as a function of flow velocities in carotid artery models of 1:1, 1:2, 1:4, and 1:6 scales after velocity adjustment using CFD analysis with (D) blood and (E) cell culture medium fluid properties. (F-G) CFD simulation derived velocity color maps at (F) physiological (1:1) and (G) 1:6 scale models. (H-I) CFD simulation derived WSS distributions at (H) physiological (1:1) and (I) 1:6 scale models.

A computer-aided design (CAD) model of a human carotid artery was first generated from clinical magnetic resonance imaging (MRI) scans from a healthy donor (Figure 1B-C). A similitude analysis to preserve the hemodynamic profile in the miniaturized carotid artery model was then performed by assuming the common carotid artery (CCA) as a straight cylindrical section with a diameter that was equivalent to the nominal lumen diameter of the CCA. The similitude analysis was based on the Buckingham Pi theorem, which yielded a non-dimensional equation that related shear stress with fluid velocity, diameter, and viscosity (Figure S1A). This simplified relationship allowed us to determine the theoretical flow velocity necessary to maintain comparable shear stress in cylinders with diameters scaled down by half (1:2), a quarter (1:4), and a sixth (1:6) (Figure S1B). To confirm these analytical findings, CFD simulations were first conducted using a straight cylinder with a diameter equivalent to that of a physiological CCA (∼6 mm) and the 1:6 scaled model (diameter ∼1 mm) to determine if the wall shear stresses matched. The simulation showed accurate reproduction of wall shear stresses (WSS) in the straight cylinder with an error of 0.7%, where the wall shear stresses for the physiological and 1:6 scaled models were 3.38 Pa and 3.42 Pa, respectively (Figure S1C). However, when we performed CFD analysis to verify the analytical scaling in the physiological and 1:6 scaled model of the carotid artery, the wall shear stresses did not match (Figure S1D). This was likely due to oversimplification of the mathematical assumptions, such as uniform flow development and symmetry, which were not fully applicable in the complex geometry of a carotid artery.

Using the velocity predicted by the similitude analysis as an initial value, we then iteratively optimized the flow velocities in the scaled-down carotid artery model to achieve shear stress values consistent with those observed in the physiological model. These adjustments were conducted using both blood and cell culture medium viscosities, with the latter being utilized in subsequent cell culture experiments. The corrected velocities effectively replicated the WSS distribution observed in physiological carotid arteries at 1:2, 1:4, and 1:6 scales for both blood (Figure 1D) and cell culture media viscosities (Figure 1E). For subsequent experimental evaluation on the relationship between vessel geometry, hemodynamic profiles and endothelium responses, we selected the 1:6 scale carotid artery model as it was an optimal balance between 3D printer resolution, fluid pump performance, standard microfluidic fittings, and cell culture requirements.

A target average shear stress of 10 dynes cm^-^² in the CCA was employed to simulate the hemodynamic profiles typically found in the carotid artery.[33] For physiological benchmarking, a flow velocity of 0.27 m s^-1^ was set as the inlet boundary condition to achieve an average shear stress of 10 dynes cm^-^² in the proximal section of a physiological scale CCA, using blood fluid properties. To ensure compliance with experimental conditions for cell culture, it is necessary to modify the scale and fluid properties. For the 1:6 model, a flow velocity of 0.21 m s^-1^ was applied with cell medium fluid properties, resulting in comparable shear stress distributions. We then compared the CFD simulations of both the physiological 1:1 (Figure 1F, H) and 1:6 scaled (Figure 1G, I) carotid arteries to evaluate the differences between the trunk of the CCA (Zone 1) and the carotid bifurcation (Zone 2), which represented high and low shear stress regions, respectively. Minor variations in the mid-plane cross-sectional velocity and WSS were observed, with errors of 1.9% and 4.2%, respectively. In Zone 1, flow patterns exhibited consistent laminar flow, with peak velocities reaching approximately 0.38 m s^-1^ for the 1:1 scale and 0.37 m s^-1^ for the 1:6 scale. Conversely, Zone 2 showed lower velocities with recirculating flow with eddies, where maximum velocities were below 0.042 m s^-1^ for the 1:1 scale model and 0.041 m s^-1^ for the 1:6 scale model (Figure 1F-G). Global WSS in Zone 1 ranged from 9.3 to 18 dynes cm^-^² with an average of ∼13.65 dynes cm^-^² at the physiological 1:1 scale, while the 1:6 scale model ranged from 8.9 to 17 dynes cm^-^² with an average of ∼12.95 dynes cm^-^². In Zone 2, however, flow disturbances along the outer walls reduced WSS to between 0 and 4 dynes cm^-^² for both scales, suggesting a region that is prone to atherosclerosis (Figure 1H-I). The CFD results confirmed that atherosclerosis-prone zones occurred at the carotid bifurcations and demonstrated that the hemodynamic profiles could be preserved in the miniaturized 1:6 model.

### 2.2. Experimental validation of hemodynamic flow profiles using micro-PIV

Micro-Particle Image Velocimetry (Micro-PIV) experiments were conducted to validate the hemodynamic profiles in the miniaturized 1:6 scale carotid artery model. The physical model was created using DLP 3D printing with a rigid clear resin, ensuring optical clarity for effective particle tracking. Particle tracking focused on the high-velocity laminar flow region in the CCA (Zone 1) and the characteristic low-speed eddy formation regions at the carotid bifurcation (Zone 2) (Figure 2A). The 3D-printed construct was connected via tubing to a perfusion circuit to circulate fluorescent tracer particles for Micro-PIV analysis (Figure 2B). A perfusion flow rate of 9.5 ml min^-1^ was used to achieve the adjusted input velocity of 0.21 m s^-1^, which produced an average WSS of 10 dynes cm^-^² in the CCA (Zone 1). The experimental results indicated that the maximum velocities within the 3D-printed 1:6 model ranged from 0.0 to 0.35 m s^-1^, closely matching the velocity contours predicted by the CFD simulation. The velocity vector maps showed that the highest velocity occurred near the bifurcation entrance of the CCA, reaching approximately 0.35 m s^-1^ in Zone 1 (Figure 2C). This measured velocity corresponded closely to the CFD-predicted velocity of approximately 0.36 m s^-1^, resulting in an error of only 2.8% (Figure 2C).. The maximum velocities measured in the bifurcation region (Zone 2) also showed minimal deviation (1.09%) from the CFD results, with the measured velocity being approximately 0.045 m s^-1^ compared to 0.041 m s^-1^ predicted by CFD (Figure 2D). WSS were calculated based on measured velocity magnitudes within the PIV model and compared with CFD predictions. High WSS in Zone 1 ranged between 5 and 17.2 dynes cm^-^², while atherosclerotic-prone areas in Zone 2 experienced WSS below 4 dynes cm^-^².

**Figure 2.**
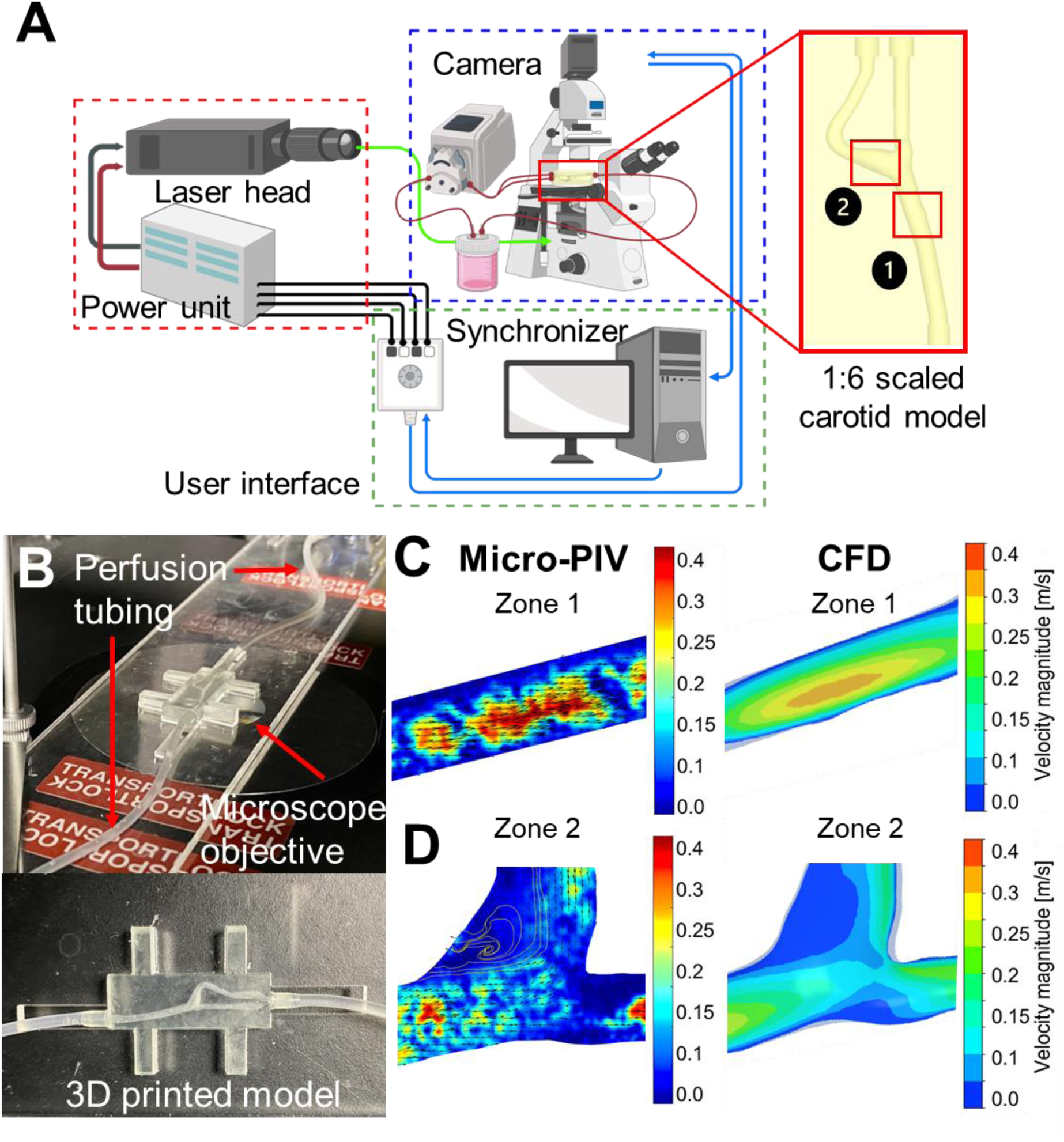
Experimental validation of hemodynamic flow profiles in a miniaturized (1:6 scale) carotid artery model using Micro-Particle Imaging Velocimetry (micro-PIV). (A) Schematic representation of the laser optics micro-PIV system setup, highlighting zones of interest: Zone 1 (common carotid artery, CCA) and Zone 2 (carotid bifurcation). (B) Images showing fluidic connection of the 3D printed carotid artery prototype to a perfusion circuit and placement on the microscope system. (C-D) Velocity profile comparisons between experimental micro-PIV results (left) and CFD simulation (right) at (C) Zone 1, corresponding to the CCA, and (D) Zone 2, corresponding to the low-speed recirculation region in the carotid bifurcation. Color gradients indicate velocity ranges (m s^-1^).

### 2.3. 3D printing and characterization of GelMA carotid artery constructs

Next, a cell-compatible, scaled-down 1:6 carotid artery model was fabricated using DLP 3D printing and a biocompatible gelatine methacrylate (GelMA) photoink. The design of the construct involved a rectangular block with the patient-derived arterial channel geometry subtracted, creating a perfusable vessel construct (Figure 3A-B). The DLP 3D printing process was highly reproducible, achieving a success rate of 90%, with 9 out of 10 constructs being printed successfully. The printed constructs exhibited excellent dimensional fidelity, with less than 5% deviation from the original CAD design in key parameters, including the overall length, width, height (Figure 3C), as well as the channel diameter and the cross-sectional areas (Figure 3D). Post-printing, we investigated how water absorption by the GelMA vessel construct impacted its dimensions after immersing it in cell culture medium for 24 hours. The results revealed that dimensional changes primarily occurred in the block’s external dimensions, with the length and width increasing by 15% and the height by 20% (Figure 3E). However, the channel inlet and outlet diameters remained largely unaffected, showing minimal swelling of about 3.5% (Figure 3E). These findings suggest that material swelling should have a limited impact during long-term perfusion culture of cells within the carotid artery construct.

**Figure 3.**
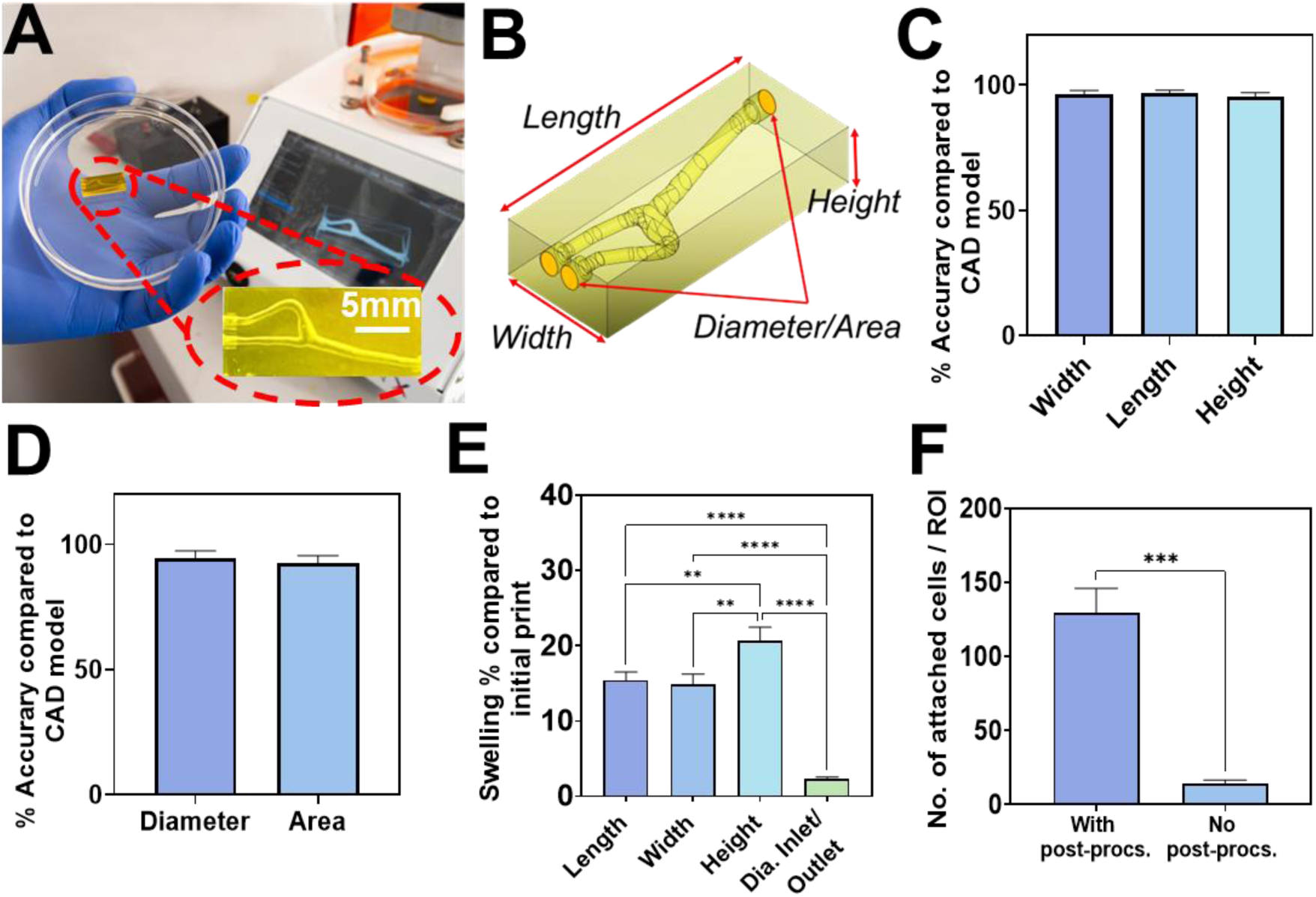
Fabrication of a DLP 3D-printed GelMA carotid artery construct. (A) Image showing the GelMA carotid artery construct after the 3D printing process. (B) CAD design of the patient-specific channel, including measured dimensions for validation. (C-D) Comparison of printing accuracy between fabricated channels and CAD models for (C) external dimensions of the entire GelMA block and (D) diameter and cross-sectional area of the perfusable vessel. (E) Dimensional changes in overall 3D printed GelMA construct and vessel diameter at the inlets/outlets after immersing in culture medium for 24 hours. (F) Effect of post-processing on hAECs attachment to 3D-printed GelMA discs 24 hours post seeding. Data are presented as mean ± SD (n=4). Statistical significance was determined using one-way ANOVA for swelling measurements and unpaired t-test for post-processing test. Statistical significance is denoted as ****p<0.0001, ***p<0.001, **p<0.01, while non-significant differences are indicated as "ns".

Since residual photoinitiators and photoabsorbers used during DLP 3D printing can exhibit cytotoxic effects, adequate post-processing is necessary to remove these compounds and ensure cell viability [34]. Here, we optimized the post-processing of 3D-printed GelMA constructs by performing at least three media changes over 24 hours. This protocol significantly enhanced the viability and attachment of seeded hAECs on the GelMA substrate compared to unprocessed substrates (Figure 3F). Despite these improvements, it is well-documented that cell adhesion and spreading on hydrogel-based substrates, such as GelMA, are inherently weaker than on tissue culture polystyrene (TCPS).[35] Our findings corroborated these observations, as we observed that a five-fold increase in cell seeding density was required to achieve comparable cell density on GelMA substrate compared to TCPS substrates (Figure S2).

Hence, to improve cell adhesion on the photo-crosslinked GelMA substrates, extracellular matrix (ECM) components were incorporated into the photo-resin. Collagen type I or fibronectin were blended with the GelMA photo-resin at a 9:1 (v/v) ratio, ensuring that the mechanical stability and 3D printing fidelity of the hydrogel vessel constructs remained uncompromised. Cell viability and density were assessed using live/dead staining on day 16 of culture. Across all conditions, cell viability remained consistently high, with no statistically significant differences observed between GelMA-only and GelMA+ECM substrates (Figure 4A-B). However, the incorporation of collagen or fibronectin markedly enhanced the number of adhered cells compared to GelMA alone (Figure 4C). This improvement in cell attachment was accompanied by an increased cell growth rate on both GelMA+collagen and GelMA+fibronectin substrates, as demonstrated by the PrestoBlue metabolic assay over a 16-day culture period (Figure 4D). Although collagen and fibronectin were comparable in promoting cell adhesion, collagen type I was selected for subsequent experiments due to its well-established role in supporting endothelial cell adhesion, proliferation, and growth. [36]

**Figure 4.**
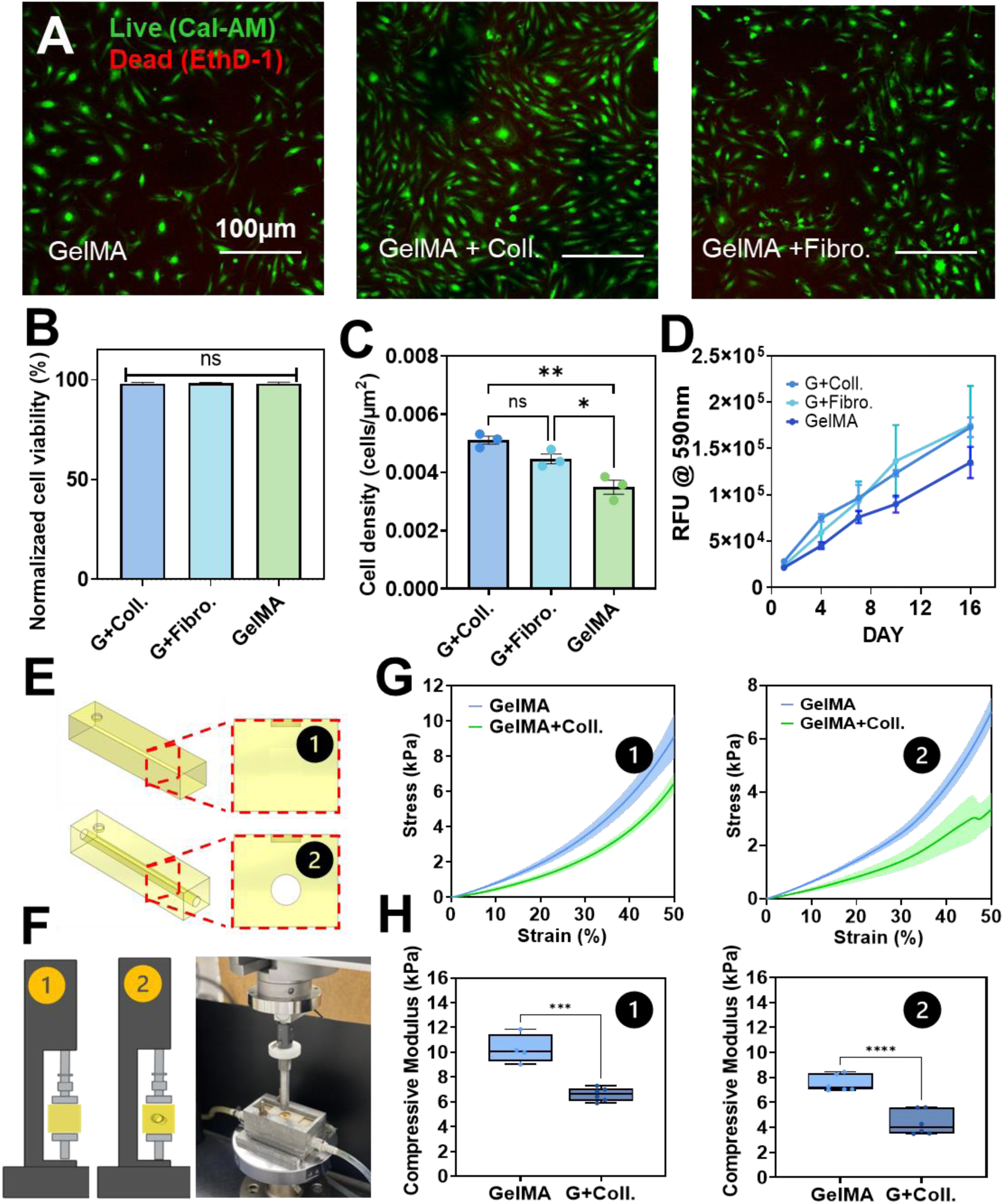
Functionalization and mechanical evaluation of DLP 3D printed GelMA substrates. (A) Live (Calcein-AM)-dead (Ethidium Homodimer-1) staining of hAECs cultured on 3D printed GelMA and GelMA+ECM (GelMA-Collagen and GelMA-Fibronectin) substrates 16 days post-seeding. (B) Quantification of cell viability for GelMA+ECM conditions compared to GelMA-only controls. (C) Quantification of density of adhered cells on GelMA+ECM and GelMA only substrates 24 hours post-seeding. (D) Cell growth rate of hAECs measured by PrestoBlue metabolic assay on GelMA+ECM and GelMA-only substrates over a 16-day culture period. (E) Schematic showing two constructs used for mechanical testing: (1) solid block and (2) block with an internal cylindrical channel of 1 mm diameter. (F) Experimental setup for compression testing using an Instron mechanical testing system. (G) Stress-strain curves and (H) compressive modulus values of GelMA and GelMA+collagen constructs, comparing (1) solid block (left) and (2) block with internal channel (right) configurations. Statistical significance was determined using one-way ANOVA for cell attachment and t-test for mechanical test. Data are presented as mean ± SD, n = 3 (cell viability / attachment) and 6 (mechanical test). Significance levels: ****p < 0.0001; ***p < 0.001; **p < 0.01; *p < 0.05; non-significant results are denoted as "ns".

We next evaluated whether the incorporation of ECM components affected the mechanical properties of vessel constructs that were 3D printed with collagen-blended GelMA (GelMA-collagen) as compared to those fabricated using GelMA alone. This was crucial to assess potential implications for their handleability and integration into a perfusion circuit. Two structural configurations were analyzed: a solid hydrogel block and a hydrogel block containing a straight cylindrical channel (Figure 4E). Compression testing was performed using an Instron mechanical testing apparatus under controlled thermal conditions in a confined chamber (Figure 4F). The stress-strain profiles demonstrated an exponential increase in stress with applied strain across both geometrical configurations and hydrogel formulations. Constructs fabricated with GelMA-collagen exhibited a slower rate of stress increase under compression, indicative of a softer and more elastic material compared to unmodified GelMA (Figure 4G). For the solid hydrogel blocks, GelMA-collagen constructs displayed a significantly lower compressive modulus (6.5 kPa) compared to GelMA-only constructs (10 kPa) (Figure 4H). For the channel-containing block, both materials exhibited reduced compression module relative to their solid counterparts. GelMA-only constructs had an average modulus of 8 kPa, which was significantly higher than the 4 kPa observed for GelMA-collagen constructs (Figure 4H). Despite the observed reduction in mechanical properties for GelMA-collagen constructs, they retained sufficient mechanical stability to withstand perfusion forces while potentially offering enhanced compatibility with cellular adhesion and proliferation. The decrease in stiffness may also improve biocompatibility, as softer materials better mimic the physiological compliance of native vascular tissues,[37] supporting their role in long-term endothelialization and functional integration within a bioreactor system.

### 2.4. Perfusion culture of hAECs and hASMCs in the 3D printed carotid artery construct

The human carotid artery consists of distinct vascular layers, each defined by specific cell types and extracellular matrix (ECM) components. In this study, we aimed to replicate the tunica intima, the endothelial cell-lined inner lumen, and the tunica media, a matrix-rich layer populated by vascular smooth muscle cells, within the 3D-printed carotid artery construct. To achieve this, we first evaluated the establishment of human arterial endothelial cells (hAECs) cultures in the scaled-down (1:6) GelMA-collagen carotid artery construct under static and dynamic conditions. hAECs were seeded into the 3D printed carotid artery construct and maintained under static culture for 10 days to assess uniformity of cell distribution, lumen coverage, and cell viability. Confocal tile-scan imaging of phalloidin-labeled hAECs confirmed uniform endothelial cell coverage throughout the lumen of the hydrogel construct (Figure 5A). Cell viability assessments using live-dead staining revealed high cell viability after 10 days of culture, which was comparable to traditional 2D culture on both TCPS and GelMA-coated plates (Figure 5B-C). There were no statistically significant differences in cell area coverage between the CCA (Zone 1) and carotid bifurcation (Zone 2) regions within the vessel construct and the 2D control groups (Figure 5D).

**Figure 5.**
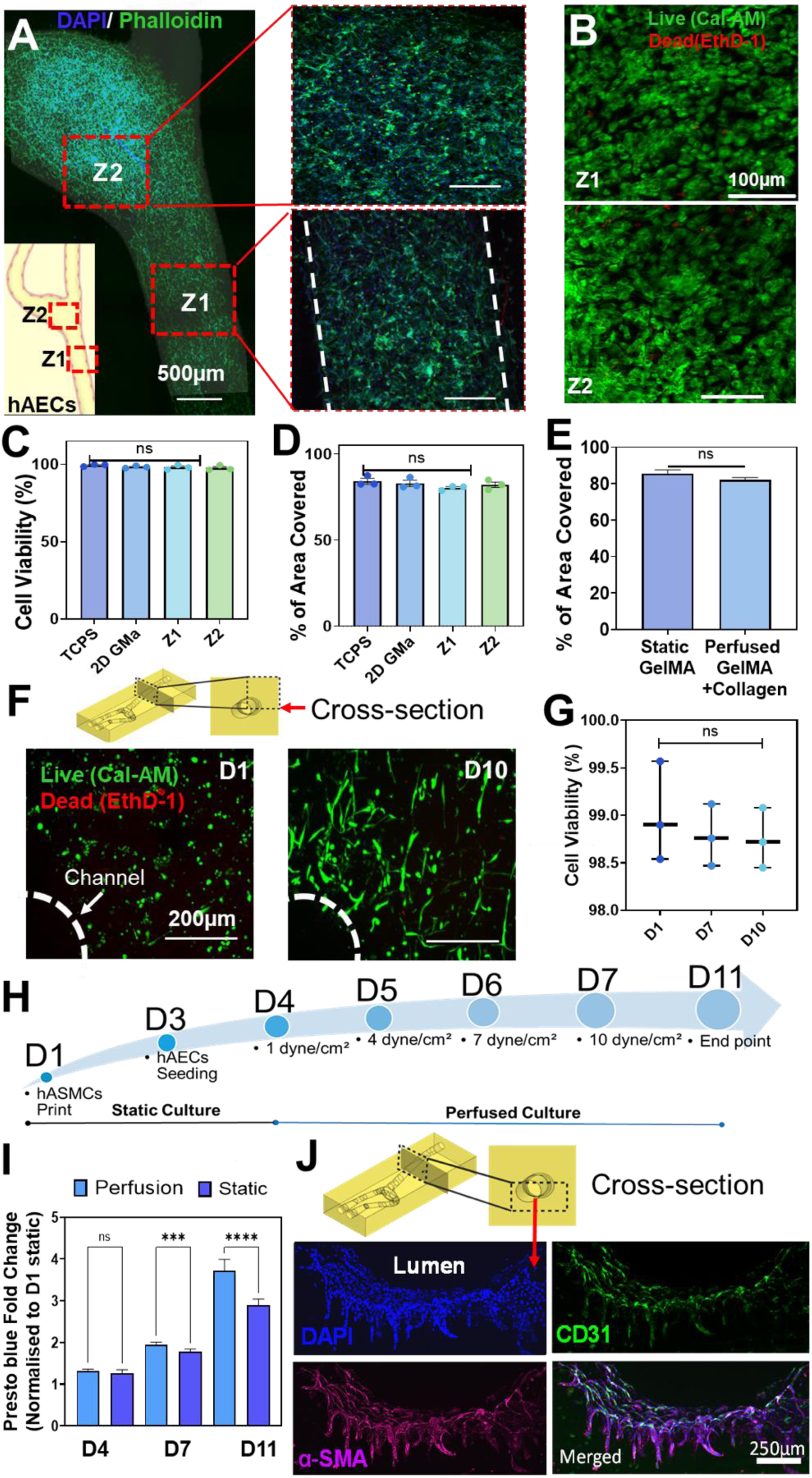
Vascular cell culture within 3D printed GelMA carotid artery constructs. (A) Confocal tile scan of the vessel construct, highlighting the common carotid trunk (zone 1) and the carotid bifurcation (zone 2) after 10 days of static culture. (B) Live-dead staining of hAECs at zone 1 (Z1, top) and zone 2 (Z2, bottom) after 10 days of static culture. (C-D) Quantification of hAECs (C) cell viability and (D) area coverage across zone 1 and zone 2 of GelMA vessels compared to those cultured on TCPS and 2D GelMA cultures. (E) Percentage area coverage of hAECs on GelMA-collagen vessel construct under perfusion culture compared to static culture on GelMA alone. (F) Live-dead staining of embedded hASMCs at days 1, and 10. (G) Quantitative assessment of hASMCs viability at days 1, 7, and 10 of culture. (H) Timeline showing seeding and perfusion culture of hAECs and hASMCs co-culture. (I) Presto Blue metabolic activity assay comparing static versus perfused co-cultures within GelMA-collagen vessel constructs over a period of 11 days. (J) Immunofluorescence images of cross-sections from co-cultured vessel constructs showing nuclei (DAPI, blue), hAECs (CD31, green), and hASMCs (α-SMA, purple). Images were acquired at 10x magnification (scale bar = 250 µm). Statistical analysis was performed using one-way ANOVA. Data are presented as mean ± SD (n = 3); statistical significance is denoted as ****p<0.0001, **p<0.001, **p<0.01, with ns indicating non-significance.

To evaluate the integrity and attachment of the endothelial monolayer within the hydrogel channels under dynamic flow conditions, we systematically tested varying wall shear stresses. The flow rates were adjusted to achieve wall shear stresses of 1, 4, 7, and 10 dynes cm^-^² in the CCA region of the construct. The actual flow rates in the vessel constructs were validated using an in-line flow sensor integrated into the perfusion loop, which exhibited less than 5% deviation from the set flow rates, ensuring precise and consistent experimental conditions (Figure S3A-B). Next, hAECs were seeded into the vascular constructs and allowed to adhere under static culture conditions for 24 hours before being subjected to dynamic perfusion. The shear stress was incrementally increased from 1 to 10 dynes cm^-^² over 5 days to mimic physiological arterial conditions (Figure S3C). Quantitative analysis of the percentage of cell area coverage in the high-velocity CCA region (Zone 1) revealed no significant differences between static and dynamic culture conditions (Figure 5E). These findings indicate that hAECs were able to attach and maintain a confluent monolayer on the lumen of the carotid artery construct under both static and perfusion culture conditions.

To replicate the tunica media layer of the vascular wall, human arterial smooth muscle cells (hASMCs) were embedded directly into the GelMA-collagen resin during DLP 3D printing. The inclusion of hASMCs has been reported to play a critical role in supporting the long-term maintenance of the endothelial layer by preventing delamination and enhancing vessel functionality in engineered constructs.[29] Live/dead microscopy imaging of orthogonal channel sections was conducted at multiple time points (days 1, 7, and 10) to evaluate cell viability and spatial distribution (Figure 5F). The results demonstrated that over 98% of hASMCs remained viable by day 10. Importantly, there were no significant changes in viability between day 1 and day 10 (Figure 5G), indicating that the DLP 3D printing process and subsequent perfusion culture did not induce cytotoxicity or impair cell survival. Moreover, by day 10, the hASMCs exhibited morphological characteristics consistent with their physiological phenotype, including elongated cell shapes and strategic positioning near the perfused channel. This adaptation suggests that the 3D microenvironment provided by the GelMA-collagen construct was conducive to proper cellular function (Figure 5F).

A custom 3D-printed perfusion bioreactor system was developed to enable dynamic, long-term co-culture of hASMCs and hAECs within the carotid artery construct by providing controlled luminal flow for endothelial cells and simultaneous support for smooth muscle cells in a dual-medium environment (Figure S4). The sequential seeding and culture of the two vascular cell types is shown in Figure 5H. hASMCs were mixed, suspended in the GelMA-collagen photoresin and 3D printed within the patient-specific carotid artery hydrogel constructs. The embedded hASMCs were maintained in static culture for 3 days before hAECs were seeded into the channel lumen. The seeded vessel constructs were maintained in static conditions for another 24 hours to allow the endothelial cells to attach. Perfusion culture was initiated on day 4 at low shear stress (1 dyne cm^-^²), and the flow rate was gradually stepped up to reach final shear stress of 10 dynes cm^-^² by day 7. This gradual increase in flow rates ensured that the hAECs could form a stable endothelial monolayer without delaminating under high-stress stress.

To continuously monitor the metabolic activity of vascular cells within the 3D-printed carotid artery constructs, PrestoBlue metabolic assay was performed over an 11-day culture period. A significant increase in metabolic activity was observed in the perfused co-cultures compared to static controls, with the difference becoming significantly pronounced from day 7 to day 11 (Figure 5I). This increase suggests enhanced cell viability and proliferation, likely due to improved nutrient delivery and waste removal under dynamic flow conditions. Following 7 days of perfusion, the constructs were fixed, cross-sectioned at the CCA region (Zone 1), and analyzed via immunofluorescence staining to assess the spatial distribution of hAECs and hASMCs. Immunostaining confirmed the presence of endothelial marker CD31 within the vessel lumen, while smooth muscle actin (SMA) was localized in the surrounding hydrogel matrix, verifying the compartmentalization of the two cell types (Figure 5J). Notably, endothelial cells exhibited infiltration into the GelMA hydrogel along the 3D printing layers, suggesting active cellular remodeling in response to the engineered vessel structure and perfusion conditions.

### 2.5. Recapitulating differential vascular cell responses to anatomically dependent hemodynamic flow in the carotid artery

Finally, we demonstrated the utility of the 3D-printed miniaturized patient-specific carotid artery model in recapitulating the interplay between anatomical-dependent hemodynamic changes and vascular cell responses. Vascular endothelial cells are highly mechanosensitive, adapting their morphology and function to the local fluid shear stress environment.[38, 39] Based on our CFD and micro-PIV results (Figure 1 and 2), the carotid artery construct exhibited two distinct hemodynamic regions: the CCA trunk, where flow was unidirectional and associated with high wall shear stress (∼12.95 dynes cm^-^²), and the carotid bifurcation, where flow was slower, recirculating, and characterized by low wall shear stress (<0.4 dynes cm^-^²). Given these differences, we examined endothelial cell alignment as a functional response to local shear stress variations after seven days of perfusion culture, including four days at the target shear stress of 10 dynes cm^-^² (Figure 6A-B). Quantitative analysis revealed significant differences in endothelial cell orientation between the two regions. In the CCA (Z1), approximately 75% of cells exhibited alignment with the direction of flow, whereas at the carotid bifurcation (Z2), alignment was reduced to ∼50% (Figure 6C). Further analysis of nuclear orientation in the CCA showed a preferential alignment, with ∼45% of cells falling within the first angle segment (68.5°–90°) and ∼35% within the second segment (46°–67.5°) (Figure 6D). In contrast, cells in the bifurcation region displayed a more dispersed alignment pattern, with no single segment exceeding 30%, indicating a loss of directional preference due to the disturbed flow conditions at the bifurcation (Figure 6E).

**Figure 6.**
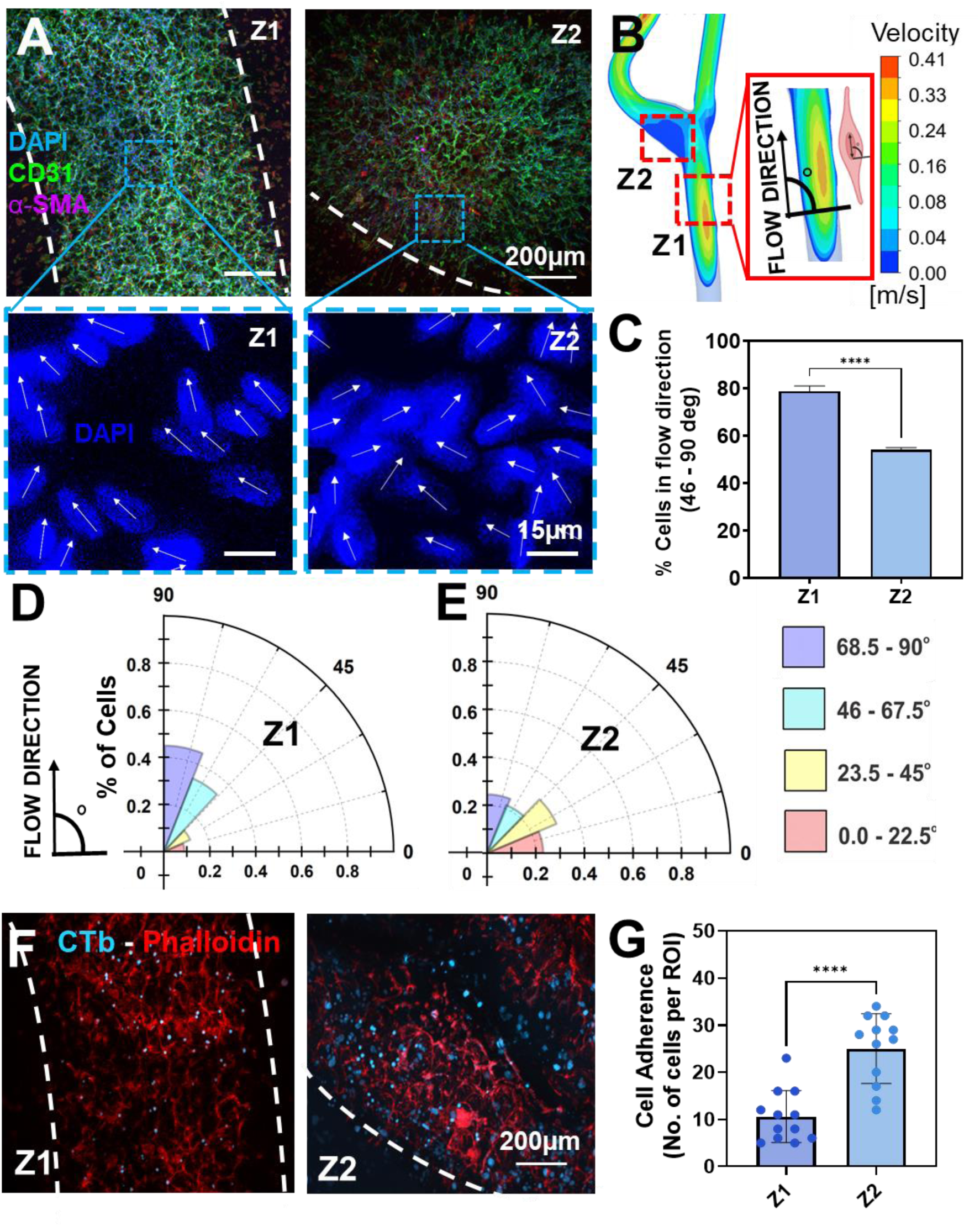
Regional differences in cell alignment and monocyte adhesion in 3D-printed carotid artery constructs. (A) Immunofluorescence of hAECs (CD31, green) and hASMCs (α-SMA, purple) in the CCA (zone 1) and carotid bifurcation (zone 2) after 3 days of static and 8 days of perfusion culture. Nuclei stained with DAPI (blue). (B) Schematic showing flow direction and cell nuclei orientation; cell alignment defined as 45°–90° to the horizontal. (C) Quantification of % hAECs aligned to flow in each zone. (D–E) Polar plots showing endothelial alignment across four angular bins. (F) Monocyte (U937, blue) adhesion on inflamed hAECs (red, Phalloidin) in zones 1 and 2. (G) Quantification of monocyte adhesion post-TNF stimulation. Data shown as mean ± SD (n = 4); ****p < 0.0001, Student’s t-test.

Endothelial cell misalignment due to hemodynamic disturbances, particularly at the carotid bifurcation, is strongly associated with endothelial dysfunction and an increased risk of atherosclerotic plaque formation.[40, 41] Dysfunctional endothelial cells exhibit a pro-inflammatory phenotype, characterized by the upregulation of endothelial-leukocyte adhesion molecules such as E-selectin and VCAM-1, which facilitate monocyte adhesion and contribute to atherogenesis.[42–44] To examine the effects of anatomical-dependent hemodynamic variations on endothelial inflammatory responses, we assessed monocyte adhesion to hAECs under inflammatory conditions. A previously established 2D monocyte adhesion assay [45] was adapted for our perfusion circuit setup by first optimizing the TNF-α concentration required to induce inflammation. hAECs monolayers cultured in 24-well plates were treated with increasing TNF-α concentrations (0, 1, 5, 10, 20, and 40 ng mL^-1^) (Figure S5A). Monocyte adhesion significantly increased with TNF-α treatment, peaking between 5 and 20 ng mL^-1^ (Figure S5B). However, at 40 ng mL^-1^, monocyte adhesion declined, likely due to TNF-α-induced cytotoxicity, as previously reported.[46] Based on these results and prior studies, 10 ng mL^-1^ TNF-α was selected for subsequent experiments.

The monocyte adhesion assay was conducted in the 3D-printed GelMA-collagen carotid artery constructs, which had been seeded with hAECs and hASMCs and maintained under optimized co-culture perfusion conditions for seven days, including four days at a target shear stress of 10 dynes cm^-^². Following this period, the vessel constructs were perfused with 10 ng mL^-1^ TNF-α for 5 hours to induce endothelial inflammation, after which fluorescently labelled U937 monocytes were introduced to assess adhesion. Monocyte adhesion was significantly higher at the carotid bifurcation (zone 2) compared to the CCA (zone 1) (Figure 6F-G). In all, our findings align with previous studies using microfluidic devices that demonstrated the correlation between endothelial alignment, inflammatory responses, and shear stress directionality.[47, 48] However, rather than relying on simplified geometries or separate channels with fixed shear stress, our patient-specific carotid artery model enabled the capture of localized variations in hemodynamic within a single construct derived from clinical imaging data.

## 3. Discussion

Recapitulating the complex interplay between vascular anatomy, hemodynamic, and cellular responses remains a major challenge for existing vascular models. While substantial progress has been made in reconstructing patient-specific anatomy, computationally predicting hemodynamic parameters, and engineering simplified biological microenvironments, most models fail to integrate all three critical aspects of vascular homeostasis, limiting their physiological relevance.[49, 50] Recent advances in 3D printing of cellularized constructs derived from clinical imaging have demonstrated potential in replicating biomimetic vascular microenvironments.[30–32] However, they have yet to demonstrate that local anatomically dependent hemodynamic flows can influence endothelial remodeling. This study addresses this limitation by developing a patient-specific carotid artery model to experimentally validate the influence of hemodynamic forces on endothelial function and their role in CVD pathophysiology. By integrating clinical imaging, computational simulation and DLP 3D bioprinting, we could translate MRI-derived hemodynamic flow data into a dynamic cell microenvironment, capturing essential aspects of vascular physiology (Figure 1A). High-resolution segmentation preserved the intricate geometries of the CCA and bifurcation regions of the carotid artery (Figures 1B-C), which is important since minor anatomical variations significantly impact local shear stress and flow dynamics.[51] The resulting anatomically accurate carotid artery CAD model facilitated detailed CFD analyses (Figures 1H-K), which were subsequently translated into physical models for micro-PIV flow validation (Figure 2B) and a bioprinted vessel model that can be cellularized (Figure 3A).

Miniaturization of the carotid artery model to 1:6 was necessary to balance fabrication feasibility, material consumption, and reproducibility of the 3D bioprinted, cell-laden constructs. While miniaturization often compromises anatomical fidelity,[52] our approach preserved key geometrical features using CAD-based scaling while ensuring hemodynamic relevance through analytical similitude analysis.[53] Iterative tuning of the flow velocity in the scaled-down models using CFD simulations corrected discrepancies in analytical scaling assumptions, aligning the shear stress distribution within the 1:6 scale model with that of the 1:1 physiological model (Figures 1H-K, Supplementary Figure 1C). Experimental validation using micro-PIV confirmed that the physiological 1:1 and 1:6 scaled-down models exhibited distinct flow characteristics of the carotid artery. High shear stress in unidirectional flow regions in the CCA (Zone 1) and low shear stress with recirculation in disturbed flow regions in the carotid bifurcation (Zone 2) (Figures 1H, 1J) align with findings linking shear stress heterogeneity to endothelial regulation.[54, 55] Low-shear regions, historically associated with atherosclerotic plaque formation, reinforce the causal relationship between disturbed flow and disease progression.[56, 57]

Leveraging DLP 3D printing to fabricate complex geometries with commercial GelMA hydrogel bioinks, we successfully replicated the 1:6 scale patient-specific carotid artery construct. We achieved printing fidelity with deviations below 5% (Figures 3A-D), which was similar to reported specifications from the manufacturers of the GelMA bioink. Hydrogel swelling behavior informed bioreactor design, ensuring structural integrity for perfusion experiments (Figure 3E). Although GelMA bioinks are purportedly formulated for cell printing using biocompatible photoinitiators and photoabsorbers,[58] residual crosslinking agents can still have cytotoxic effects.[59] Our results demonstrated significantly higher cell attachment and viability in post-processed constructs (Figure 3F), aligning with previous studies emphasizing the importance of thorough washing steps to remove unreacted photoinitiators. Adhesion of hAECs to the 3D printed GelMA vessel construct can be further enhanced by adding ECM proteins found within the arterial wall, such as Collagen type I and fibronectin to the GelMA bioink (Figure 4A-D). However, the addition of collagen softened the hydrogel matrix, reducing stiffness by ∼4 kPa (Figure 4E-H), the resulting mechanical properties remained within the range of normotensive arterial tissue stiffness.[60, 61] This trade-off between mechanical stability and enhanced cellular interactions supports the functionalization of hydrogels for vascular applications.

Our 3D-printed carotid artery model was maintained as a perfused co-culture system, enabling the simultaneous culture of hASMCs and hAECs in separate media using a custom bioreactor (Figure S4). This dual-media approach contrasts with conventional vascular co-cultures that rely on a single medium, which may not adequately support the specific requirements of each cell type.[29] Both hASMCs and hAECs exhibited good viability under static conditions for up to 10 days (Figures 5A and 5F), likely supported by the porous nature of the GelMA hydrogel, [62] which promotes nutrient and oxygen diffusion. This porosity facilitated sequential seeding of hASMCs followed by hAECs as well as ensuring that cells were confluent prior to initiating perfusion to minimize the risk of cell detachment. Upon the introduction of flow, hAEC proliferation was further enhanced in response to shear stress (Figure S3).[63] Interestingly, we observed endothelial cell infiltration into the GelMA matrix from the vessel lumen (Figure 5I), potentially driven by shear stress gradients or migration along the layered architecture of the 3D-printed construct. The visible step-like features of the lumen, corresponding to the 10 µm printing resolution, may contribute to this behavior. Employing higher-resolution DLP printers (e.g., 2 µm) may help reduce these surface irregularities and improve barrier fidelity.

Our 3D-printed carotid artery model offers a robust and physiologically relevant platform to investigate the mechanobiological mechanisms that link local hemodynamic forces to atherosclerosis. Regional variation in endothelial responses was evident, with approximately 75% of hAECs aligning with the flow direction in the CCA (Zone 1), while cells at the carotid bifurcation (Zone 2) displayed misalignment and disorganized morphology (Figure 6C, E). This disrupted alignment is a hallmark of endothelial dysfunction, a known precursor to atherogenesis, and is often accompanied by elevated inflammatory signaling and lipid accumulation.[56] Importantly, the model also demonstrated that TNF-α-induced inflammatory responses were amplified under disturbed flow conditions at the carotid bifurcation, as evidenced by significantly increased monocyte adhesion (Figure 6F–G), in agreement with previous reports.[64, 65] These findings underscore the critical role of vessel geometry in modulating local flow and vascular inflammation. Specifically, anatomical parameters such as bifurcation angle and the internal carotid artery (ICA)/CCA diameter ratio are recognized contributors to atherosclerotic risk.[66] By enabling the recreation of patient-specific vascular geometries, our model provides a unique opportunity to study how anatomical variation influences susceptibility to atherosclerosis through its impact on local flow dynamics. This capability positions the platform as a valuable tool for uncovering mechanobiological links between vascular anatomy and disease initiation, ultimately supporting more personalized approaches to cardiovascular risk assessment and therapeutic development.[67]

## 4. Conclusions

This study presents a scalable and physiologically relevant in vitro model for investigating endothelial responses under patient-specific hemodynamic conditions. By integrating CFD simulations, micro-PIV validation, and perfusion cell culture within a 3D-printed hydrogel carotid artery construct, we offer a comprehensive platform to study the complex interplay between anatomically driven hemodynamic forces and endothelial function in vascular pathology. This paper provides a foundation for future work to incorporate pulsatile flow profiles and optimize the mechanical properties of the hydrogel bioink to more accurately replicate biomechanical fluid-structure interactions (FSI) and their influence on endothelium remodeling. These ongoing developments will further enhance the model’s translational relevance for disease modelling and therapeutic testing, bridging the gap between computational predictions and clinically meaningful in vitro experimentation.

## 5. Experimental Section

### Generation of 3D models of patient-specific carotid artery models

This study received approval from the Human Research Ethics Committee at Princess Alexandra Hospital (PAH) in Brisbane, Australia, and the Queensland University of Technology’s (QUT) Office of Research Ethics and Integrity (HREC/17/QPAH/181). Magnetic Resonance Imaging (MRI) scans from the healthy volunteers formed the basis for reconstructing the anatomical structure of the common carotid artery (CCA) and its bifurcation into the external (ECA) and internal carotid arteries (ICA). Participants were scanned using a 64-channel head/neck coil on a 3T Magnetom Prisma MR whole-body system (Siemens Medical Solutions, Malvern, PA, USA). Time-of-flight (TOF) magnetic resonance angiograms (MRA) were extracted from the multi-sequence MRI data. The scans were imported with the following parameters: a repetition time (TR) of 21 ms, an echo time (TE) of 3.11 ms, a resolution of 384x290x136 pixels, and a field of view (FOV) of 151x199 mm. The datasets were segmented and reconstructed (Amira version 6.0, FEI, Hillsboro, Oregon, USA). To enhance image visibility for precise selection of the carotid artery cross-section (lumen), pixel intensity adjustment and contrast enhancement tools were applied (Figure 1B). A semi-automated segmentation was then performed across the stack of cross-sectional images to generate a 3D reconstruction of the carotid artery. The automated selection process was reviewed to rectify range-based errors, ensuring accuracy and consistency in the reconstruction (Figure 1C). The final geometry was exported as a Standard Triangle Language (STL) mesh file. The STL carotid geometry was imported into ANSYS Workbench version 19.0 (ANSYS, Canonsburg, PA, USA). Using the SpaceClaim ANSYS module, the STL file was translated into a stitched surface volume, which was then enclosed and transformed into a computer-aided design (CAD) solid feature. The resulting solid underwent smoothing and geometrical adjustments, including length corrections and the elimination of any downstream arterial branches captured during the segmentation process. Additionally, slight extensions were applied to outlets to prevent regions of interest from being too close to CFD boundary conditions.

### Analytical similitude analysis for hemodynamic scaling

The 3D coronary arterial models were miniaturized to different extents (1:2, 1:4, 1:6) from the physiological scale. To determine the corresponding flow velocities that the miniaturized models would need to be operating at physiological wall shear stress levels, we first analytically calculated a flow velocity to produce arterial shear stress for a simplified cylindrical representation of the common carotid artery (CCA) at its physiological diameter. Next, equivalent velocities that produce corresponding shear stress to the CCA for the scaled versions were calculated analytically as briefly described. A simplified non-dimensional mathematical expression was derived through similitude analysis using the Buckingham Pi theorem, as demonstrated in Equation 1. The non-dimensional equation was calculated with shear stress as a function of fluid velocity, diameter, viscosity, and density, as presented in Equation 2. The resulting π1 term established a relationship between the two scenarios with identical variables – physiological and scaled and considering the shear stress (τ) (Equation 3). The resulting scaled velocity values were used as boundary conditions in CFD simulations in equivalent cylindrical geometries and then applied to simulations on the reconstructed arteries at different scales.

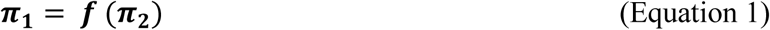

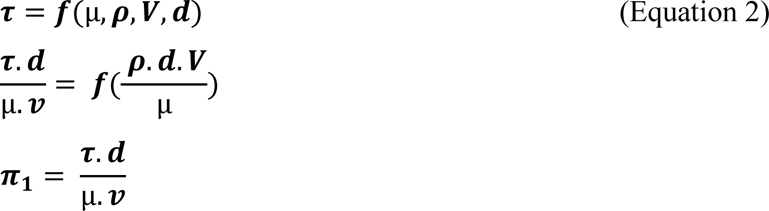

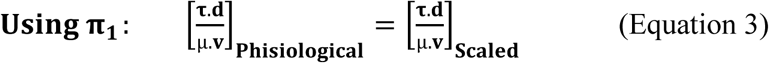

### Computational fluid dynamic (CFD) simulations on physiological and scaled-down carotid artery models

Post-processed CAD geometries were imported into the Fluent module of ANSYS Workbench version 19.0 (ANSYS, Canonsburg, PA, USA) to examine hemodynamic flow within the bifurcated carotid artery geometries. Fluid dynamic simulations were executed at both physiological and downscaled scale ratios of 1:2, 1:4 and 1:6.

The geometry meshing protocol for simulations was based on an element size independence test to ensure robustness. For the physiological model, a mesh configuration with an element size of 0.4 mm (comprising 1,148,430 elements) was chosen. For the scaled models, an element size of 0.2 mm, 0.1mm, and 0.07 mm (consisting of 813,145, 738,738, and 673,382 elements, respectively) for the scales 1:2, 1:4, and 1:6 were selected, respectively. Fluid properties were defined with laminar regime characteristics and a rigid no-slip boundary condition. The fluid was modelled as an incompressible Newtonian fluid, with viscosity values of 0.00345 Pa*s (blood)[68] for physiological conditions and 0.000825 Pa*s (cell media) [69] for scaled conditions. To mimic physiological shear stress conditions in arteries, a flow velocity was analytically calculated and used as a boundary condition to generate approximately 10 dynes cm^-^² within the CCA section at the physiological scale. A velocity magnitude of 0.272 m s^-1^ with blood viscosity was used for the 1:1 scale, while 0.42, 0.275, and 0.21 m s^-1^ were used respectively for the scales 1:2, 1:4, and 1:6 using cell media viscosity after a systematic velocity increment tuning from the initial analytical value. Outlet boundary conditions were set with 60% outflow directed to the external carotid artery (ECA) and 40% to the internal carotid artery (ICA) for both cases [68].

### Micro-particle image velocimetry (PIV) analysis

Physical models of the 1:6 scaled-down carotid artery were 3D printed to experimentally determine flow profiles with micro-PIV. The CAD models were designed (SOLIDWORKS 2021, Dassault Systems, France) to include barbed ends at the inlets and outlets for tubing connection, allowing for perfusion. The STL versions of the CAD models were sliced using ASIGA Composer slicing software (Version 1.3, ASIGA), and then 3D printed using the Asiga Max X27 DLP 3D printer with a 385 nm LED light source (ASIGA, Australia) with Moiin TechClear resin (DMG Digital Enterprises SE, Germany). The printing was done on a glass surface building plate to ensure bottom transparency.^77^ Post-printing, the 3D printed constructs underwent a 10-minute wash in isopropyl alcohol (IPA, Sigma-Aldrich, Australia), followed by a 2-hour UV curing process.

To drive fluid through the 3D-printed constructs, a peristaltic pump (IP 65, ISMATEC) was used. Flow control and isolation were ensured through switch valves placed before and after the 3D-printed device. Falcon tubes, equipped with tubing adapter lids, functioned as reservoirs for the circuit. Silicon tubing was used between the various segments of the circuit with barbed connectors. The peristaltic pump operated at a flow rate of 9.8 mL min^-1^, corresponding to a velocity sufficient to induce approximately 10 dynes cm^-^² on the wall of the scaled model, as determined by adjusted dimensional analysis calculations for the microfluidic scale.

The Micro-PIV system employed a DualPower laser (100/200 Hz, Dantec Lasers) that was transmitted through fiber optics wire to the microscope objective. The laser beam was focused onto the circular plane visualized by the microscope, which cross-sectioned the 3D-printed carotid model. For optimal image clarity, fluorescent particles (PMMA-RhB-Frak-Particles, DANTEC, Germany) in a 25% w v^-1^ aqueous suspension were employed. These particles, ranging in size from 1 to 20 µm, were excited at 560 nm to emit light at 584 nm. The emitted light was captured using a fluorescence filter, effectively capturing the specific wavelength emitted from the particles, minimizing noise from light refraction and external reflections. A high-speed camera (FlowSense EO 5M, 2448 x 2058 px, DANTEC Dynamics, Denmark) was synchronized with the laser source using a performance synchronizer (DANTEC Dynamics, Denmark) to capture the fluorescent particles flowing through the channel. A 2.5x magnification objective was used to track particle motion, while a long-pass filter (540 nm) isolated the emissions from the fluorescent particles. Data acquisition included three sets of images, each containing 10 pairs of images captured at a 10 Hz trigger rate with 1500 µs intervals between pulses. Post-processing of the particle images was conducted using MATLAB (Natick, Massachusetts, USA) with the PIVLab plugin. This included background noise removal, image masking, and the application of a multiphase cross-correlation algorithm. The interrogation window size was set at 64 × 64 pixels, with amplification up to 128 × 128 pixels to enhance correlation accuracy.

### Preparation of photo-resins

Gelatine Methacrylate (GelMA) PhotoInk™ (CELLINK, Switzerland) served as the primary photo-resin used to generate hydrogel carotid artery constructs. The addition of natural extracellular matrix (ECM) components was used to enhance cell adhesion using a formulation of 9 parts GelMA photo-resin base to 1 part ECM. The ECM components, including Collagen type I (PureCol 3 mg ml^-1^, C #5005, Advanced Biomatrix, USA) and Fibronectin (Fibronectin human plasma 1 mg ml^-1^, F1056-1MG, Sigma Aldrich, Australia), were prepared separately from their stock solutions, ensuring final concentrations of 0.3 mg ml^-1^ and 0.1 mg ml^-1^, respectively. The GelMA-ECM photo-resins were gently mixed using a piston pipette and then heated to 37°C for 10 minutes in a water bath. Upon reaching the desired temperature, the photo-resin was either directly dispensed onto the 3D printer or mixed with a cell suspension before dispensing onto the printer.

### 3D Digital Light Processing (DLP) bio-printing of carotid artery constructs

Carotid channels scaled 1:6 times were 3D printed using the Lumen X DLP 3D printer (CELLINK, Switzerland) with GelMA PhotoInk™ photo resin, both alone and in combination with ECM (collagen type I). The models were sliced into 50 µm layers with a 5.5-second exposure for each layer and a 3x exposure time for the burn-in layer at 20 mW cm^-^² projector light intensity. The hydrogel constructs were gently removed from the building platform using a plastic razor and placed in a petri dish or well plate. The channels were then internally flushed three times with 1× Phosphate buffered saline (PBS) or cell culture media to remove any uncured hydrogel. Following this, the constructs were immersed in fresh cell culture media. For printing characterization, the channels were stored at 4°C and maintained with fresh media for 24 hours to thoroughly wash the bulk material and eliminate any residual photo-initiators or photo-absorbers.

### Accuracy and swelling assessments on 3D printed GelMA carotid artery constructs

Bright-field microscopy (Stereomicroscope, Nikon SNZ1500, Japan) was utilized to evaluate the dimensions of the 3D-printed carotid hydrogel constructs (n=4) both immediately post-fabrication and following a 24-hour immersion in cell culture media. Images were captured at 1.6× magnification and measurements from the outer cuboidal structure, such as height, length, and width were measured using ImageJ software (FIJI, USA). Additionally, the diameter and area of the channel inlets and outlets were also measured. Upon removal from the cell media, the constructs were carefully dried with Kimwipes and subsequently re-measured. Dimensional deviations from the original CAD model were quantified to calculate the percentage of error, assessing the accuracy and stability of the constructs over time. Additionally, by comparing the dimensions recorded after 24 hours to those taken immediately post-printing, the percentage of swelling was determined. These data are critical for informing bioreactor design, considering the hydrogel’s dimensional changes over time.

### Mechanical testing of 3D printed GelMA carotid artery constructs

The mechanical properties of the DLP 3D printed carotid hydrogel constructs were assessed through unconfined compression tests using the Instron 68TM-30 (Instron Corp, Norwood, Massachusetts, USA). For this evaluation, cubic sections (3.5 × 3.5 × 3.5 mm³) were printed with GelMA and GelMA-Collagen, both with and without an internal channel. The samples were incubated in cell culture medium overnight to reach their fully swollen dimensions. Mechanical tests were carried out at 37°C, using a water bath connected to the compression chamber to keep the samples hydrated through the measurement process. The unconfined compression test was conducted at a displacement rate of 10 μm s^-1^, subjecting the samples to 50% compression. The stress vs. strain curve was obtained using the following mathematical relationships:

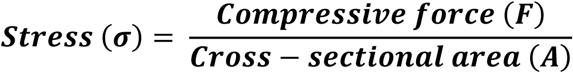

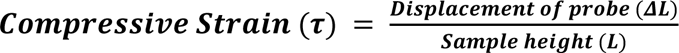 these equations, σ represents stress, F denotes the compressive force, A is the sample’s cross-sectional area, τ represents compressive strain, L stands for the sample height, and *ΔL* quantifies the displacement of the probe. The compressive modulus was derived by performing linear regression through the data points extracted from the stress vs. strain curve. A total of six replicates were tested (n = 6).

where, σ represents stress, F denotes the compressive force, A is the sample’s cross-sectional area, τ represents compressive strain, L stands for the sample height, and *ΔL* quantifies the displacement of the probe. The compressive modulus was derived by performing linear regression through the data points extracted from the stress vs. strain curve. A total of six replicates were tested (n = 6).

### Design and fabrication of perfusion loop and bioreactor

To enable continuous perfusion of the 3D printed hydrogel artery constructs, a custom enclosure chamber and bioreactor were designed using CAD software (SOLIDWORKS 2021, Dassault Systems). The bioreactor featured a support frame with one inlet and two outlets. The bioreactor can be fitted around the hydrogel artery constructs such that the lumen is connected via internal fittings and barbed connectors for external tubing. An enclosure chamber was then designed to submerge the bioreactor containing the hydrogel artery construct in cell culture medium. This design allowed cells within the 3D printed hydrogel artery construct to be fed from both the abluminal and luminal sides by culture medium contained within the enclosure chamber and medium perfused through the vessel construct, respectively. The enclosure chamber featured perforations on each side to allow tubing connection to the bioreactor and a glass slide bottom to enable visualization of the vessel construct over the culture period.

Both the bioreactor and enclosure chamber were fabricated using stereolithography (SLA) 3D printing (SLA 3+, FormLabs, Somerville, Massachusetts, USA) using a 405 nm light source. The STL files from the CAD designs were uploaded to PreForm slicing software (FormLabs Ver. 3.30.0, Formlabs Inc., Somerville, Massachusetts, USA) before being printed using BioMed Clear resin (FormLabs, Somerville, Massachusetts, USA) with an adaptive layer thickness mode to ensure good printing fidelity. The printed devices then underwent post-processing via a 10-minute soak and sonication process in 98% IPA (Sigma-Aldrich, Australia), followed by a UV-light post-curing process for 2 hours to ensure the component’s structural integrity. For sterilization, the 3D-printed components were soaked in 80% ethanol for 2 hours and washed 3 times with 1× PBS before assembling the microfluidic circuit.

Perfusion experiments were conducted using a perfusion loop connected to the enclosure chamber and bioreactor, allowing continuous circulation of the cell medium and long-term perfusion cell cultures. The perfusion loop was placed on a custom-built base holder laser-cut from acrylic sheets. The setup included a 4-way valve, switch valve, the bioreactor enclosure, and a modified 50 mL Falcon tube as a reservoir. All standard microfluidic components were sourced from IDEX Health & Science LLC (Rohnert Park, California, USA). The media flow was driven by a peristaltic pump (IP65 ISMATEC, Masterflex, Gelsenkirchen, Germany).

### Cell culture of hAECs and hASMCs

Human aortic endothelial cells (hAECs) (TeloHAEC, CRL-4052™, ATCC, USA) and human aortic smooth muscle cells (hASMCs) (AoSMC, CC-2571, Lonza, Switzerland) were routinely maintained in T175 cell culture flasks at 37.5°C and 5% CO₂. hAECs were grown in Vascular Cell Basal Medium supplemented with the Endothelial Cell Growth Kit-VEGF (PCS-100-041, ATCC, USA) and hASMCs were grown in the Smooth Muscle Cell Growth Medium 2-Bullet Kit (CC-3182, Lonza, Switzerland). 1% penicillin-streptomycin (ThermoFisher, Australia) was supplemented in all media. All cells were passaged at a 1:4 ratio when they reached 90% confluency. Passage 8 or lower was used for all cell experiments in the study.

### Cell seeding and culture in 3D printed GelMA carotid artery constructs

To seed hAECs into the 3D printed artery construct, a stock cell suspension of 10 million cells mL^-1^ was prepared. The hydrogel was assembled with the bioreactor, and the cell suspension was perfused through the circuit. Approximately 250 µL (2.5 million cells/block) of cell suspension was introduced into the 3D printed hydrogel channels through the 4-way valve of the perfusion circuit using a pipette. After seeding, the cells were allowed to attach for at least 24 hours before initiating perfusion or conducting downstream experiments. Perfusion culture was initiated at 0.94 mL min^-1^ (1 dynes cm^-^²) followed by a progressive increase of medium flow rates (4.0, 6.59, and 9.89 mL min^-1^) over 4 days every 24 hours, corresponding to shear stresses of 4, 7 and 10 dynes cm^-^². Constructs were maintained at the highest perfusion flow rate until the assay.

hASMCs were embedded into the GelMA-based photoresin during the 3D printing process of the carotid artery constructs. For each print, 5 million hASMCs were suspended in 700 µL of GelMA-photoresin before the fabrication of the constructs (n=4). Following the 3D printing, the cellularized constructs were incubated at 37°C in 6-well plates, with media changes performed three times within the first 24 hours to ensure optimal cell viability. Subsequent media changes were conducted once every 24 hours. The hASMCs were maintained under static culture conditions for 3 days before seeding with hAECs. After hAECs seeding, both cell types were cultured with their respective media, which was refreshed every 24 hours.

### Evaluation of hAECs attachment

The attachment of hAECs on different GelMA-based photoinks was first evaluated in 2D cultures using 24-well plates. Briefly, 200 µL GelMA-based hydrogel (GelMA, GelMA-Collagen, GelMA-fibronectin) was pipetted into each well (n=4 per condition) and exposed to UV light at a 405 nm wavelength using the LunaCrosslinker™ (GELOMICS, Australia) for 5 minutes to achieve crosslinking. The crosslinked hydrogels were post-processed by washing 3 times with 1×PBS over 24 hours before adding 500 µL of cell media to each well. hAECs were seeded at densities of 5,000 cells cm^-^² for post-processing and ECM functionalizing experiments and at 5,000 or 25,000 cells cm^-^² for seeding density studies. To evaluate cell attachment in perfused 3D printed GelMA-collagen artery constructs, hAECs were seeded as described above, then incubated for 24 hours in static culture and perfused for 5 days as described in the cell seeding and culture in 3D bio-printed carotid hydrogel constructs protocol (n=4).

For all samples, non-adherent cells were removed following a 24-hour incubation period. The adhered cells were then fixed with 4% paraformaldehyde (PFA) for 30 minutes, followed by staining with DAPI to visualize nuclei and phalloidin to highlight cytoskeletal structures. Quantification of cell attachment was performed using ImageJ software (FIJI, USA) based on DAPI staining to count individual cells. Additionally, qualitative evaluations of cytoskeletal surface coverage, both pre- and post-perfusion, were conducted through phalloidin staining, enabling visualization of cytoskeletal organization and cell distribution across the construct surfaces.

### DAPI and phalloidin staining

To examine cell nuclei and cytoskeleton, DAPI and Phalloidin staining were performed. For both 2D and 3D cultures, cells were washed with PBS and then fixed with 4% PFA (Sigma Aldrich, Australia) at room temperature for 20 minutes. Simultaneously, DAPI (ThermoFisher, Australia) 1 mg mL^-1^ stock solution was at a ratio of 1:500, and Alexa Fluor™ 488 Phalloidin (ThermoFisher, Australia) 66 µM stock solution was diluted at a ratio of 1:400 in PBS. Both components were incubated at room temperature for 30 minutes. Subsequently, three successive washes with 1X PBS ensured the removal of any excess or unbound staining components. The cells were imaged using a confocal microscope (Nikon Confocal A1, Nikon, Japan) with 10x and 20x objectives.

### Evaluation of cell metabolic activities

For metabolic assays, PrestoBlue (ThermoFisher, Australia) was mixed with cell medium according to the manufacturer’s instructions and incubated with the cells for 1.5 hours at 37°C. After incubation, supernatants were collected and dispensed into 96-well plates (100 µL each, with n=9 technical repetitions per sample). For 2D cultures in ECM functionalization experiments, PrestoBlue solution was applied to the substrate surfaces and removed after the incubation period. These assays were conducted over five-time points spanning 16 days (n=4). For 3D cultures on hydrogel carotid channels, PrestoBlue solution was pipetted into the chamber and through the channels. Supernatants were collected over 11 days, with measurements taken at four-time points (n=4). The 550/600 absorbance ratio was measured using a CLARIOstar plate reader (BMG LABTECH, Ortenberg, Germany).

### Live-dead staining

For viability assessments, a live/dead imaging kit was employed (488/570 excitation wavelength) (ThermoFisher, Australia). Initially, a 2× stock solution containing both Live (Cal-AM) and Dead (EthD-1) staining components was prepared in equal proportions with cell media, then pipetted to the cells and incubated at room temperature for 15 minutes. In 2D cultures, the live/dead kit was directly applied to the cultured well plates and incubated for 20 minutes (200 µL per well). For 3D cultures in the cell-laden 3D printed carotid constructs, the blocks were removed from the perfusion circuit and then immersed in 2 mL Eppendorf centrifuge tubes containing the staining solution (500 µL per tube). After the incubation period, the samples were imaged using a confocal microscope (Nikon Confocal A1, Nikon, Japan). Additionally, cells were fixed and counterstained with DAPI following the protocol described above. Images of live/dead and DAPI-stained cells were analyzed using the counting particles and threshold tools in ImageJ software (FIJI, USA) for cell viability and area coverage calculations.

### Immunofluorescence staining (IFC)

Initially, the 3D-printed hydrogel carotid constructs were removed from the bioreactor and fixed by immersing them in a 2 mL Eppendorf tube filled with 4% paraformaldehyde (PFA) for 40 minutes at room temperature. Following fixation, the vascular constructs were rinsed with 1× PBS three times. The cells were then permeabilized with 1% Triton-X-100 (Sigma Aldrich, Australia) in 1× PBS for 24 hours at 4°C. Subsequently, the cells were blocked for a day at 4°C using a solution comprised of 2% BSA (Sigma Aldrich, Australia) and 1% Triton-X-100 in PBS. The immunostaining protocol for hAECs and hASMCs was conducted as follows: First, a two-day incubation was carried out with primary antibodies specific to alpha-smooth muscle actin for hASMCs (mouse; Abcam, Cambridge, United Kingdom) and CD31 (rabbit; Thermo Fisher, Australia) for hAECs at a concentration of 1:200. This incubation was performed within a solution composed of 2% BSA and 0.2% Triton-X-100 at 4°C. The vascular constructs were then subjected to a washing step with a buffer solution (0.1% Triton-X-100 in 1× PBS), repeated twice, and kept in a washing buffer on an orbital shaker at 4°C overnight. Following the washing period, a two-day incubation was executed with an anti-mouse secondary antibody (donkey anti-mouse 555 nm, Thermo Fisher, Australia) and an anti-rabbit secondary antibody (donkey anti-rabbit 488 nm, Thermo Fisher, Australia) at a concentration of 1:200 in a solution of 2% BSA and 0.2% Triton-X-100 at 4°C. Unbound secondary antibodies were removed through a washing process with buffer solution twice at room temperature, for 1 hour each, and kept in washing buffer on an orbital shaker at 4°C overnight. Subsequently, counterstaining with DAPI was performed following the protocol described in the DAPI and Phalloidin staining section. Finally, the vascular constructs were rinsed with PBS three times and immersed in RapiClear gel (SunJin Lab, Taiwan) for one day before confocal imaging. The cells were imaged using a Nikon A1R confocal microscope with 10x and 20x objectives.

### Cell alignment measurements on 3D printed GelMA carotid artery constructs

Cell alignment experiments comprised a cell maturation protocol as described in the cell seeding and culture in 3D bio-printed carotid hydrogel constructs protocol and 4 days of continuous flow rate at ∼9.8 mL min^-1^ (∼10 dynes cm^-^²) (n=4). Subsequently, the blocks were fixed and stained following the protocol described in the DAPI and Phalloidin staining section.

Cellular alignment was assessed through nuclear alignment determination, following established methods (Aubin et al., 2010). Briefly, the orientation angle of the major nuclei axes of the cells against the orthogonal cross-sectional plane of the artery was measured. The measured angles were normalized from 0 to 90 degrees and grouped into angle ranges (0 to 22.5, 23.5 to 45, 46 to 67.5, and 68.5 to 90 degrees) to compare the percentage of cells near the flow direction (46 to 90 degrees). Three images were captured from each construct at zone 1 (CCA) and zone 2 (Bifurcation to ICA and ECA) using a Nikon A1R fluorescence confocal microscope. ImageJ (FIJI, USA) was utilized for processing the microscopy images. The major axis angle of 150 cell nuclei per image was measured to quantify cell angular orientation regarding the flow direction. The resulting data for nuclear/cell orientation angles for each channel were visualized as polar maps using OriginLab (OriginLab Corporation, USA). The quantified alignment is represented as the percentages of cells present within the angle range quartiles.

### Monocyte adhesion assay

Routine maintenance of U937 cells. U937 cells were labelled with 1.25 uL stock CellTracker Blue CMAC (ThermoFisher, Australia) per 5 mL serum-free medium according to an established protocol [70]. The labelled U937 cells were counted and spun down to obtain a concentration of 5×10⁵ cells ml^-1^, then suspended in a mixture of HAEC medium and RPMI 1640 medium supplemented with 10% FBS (Thermo Fisher, Australia) at a 1:1 ratio just before adding to inflamed hAECs cultures.

Inflammation of hAECs was induced with Tumor Necrosis Factor-α (TNF-α) before the monocyte adhesion assay. The concentration of TNF-α was optimized using 2D hAECs cultures in 24-well plates, seeded at a density of 50,000 cells/well. Recombinant human TNF- α (Abcam, Cambridge, United Kingdom) was reconstituted at a stock concentration of 2 μg ml^-1^ and diluted with hAECs cell culture media to obtain concentrations of 1 ng ml^-1^, 5 ng ml^-1^, 10 ng ml^-1^, 20 ng ml^-1^, and 40 ng ml^-1^. 500 μL of TNF-α at the determined working concentrations was added, followed by a 5-hour incubation period. 1 million cells of labelled U937 cells were added to each well and incubated for 1 hour. After incubation, the wells were gently washed three times with 1× PBS to remove non-attached monocytes before replacing them with hAECs-RPMI 1640 medium and imaged with a fluorescence confocal microscope.

For monocyte adhesion assays on 3D printed carotid constructs, the hAECs monolayer within the GelMA hydrogel carotid channel was seeded and matured as described in the cell attachment on GelMA substrates section with 4 days of continuous flow rate at ∼6.8 mL min^-1^ (∼10 dynes cm^-^²) (n=4). On the 5th day of perfusion, the hAECs medium was removed from the reservoir of the perfusion circuit and replaced with 5 ml of hAECs medium pre-mixed with TNF-α (10 ng ml^-1^), and similarly, hASMCs medium from the bioreactor chamber was replaced with 5 ml pre-mixed with TNF-α (10 ng ml^-1^). These solutions were incubated and perfused through the circuit at a flow rate of 0.37 mL min^-1^ (∼0.5 dynes cm^-^²) for a 5-hour perfusion period. Subsequently, the TNF-α containing medium was removed from the perfusion circuit, and the cells were washed twice by flushing 1 ml of PBS through the culture setup. Subsequently, labelled U937 cells were resuspended in a mixture of hAEC-RPMI-1640 medium at 10×10⁶ cells ml^-1^. Approximately 2×10⁶ labelled U937 cells (∼200 μL of cell suspension) were manually infused into the lumen of each carotid hydrogel channel and incubated for 1 hour. After incubation, the hydrogel channel was flushed three times with PBS to remove unattached monocytes. The hydrogel construct was then fixed with 4% PFA for 15 minutes, followed by washing with PBS three times and Phalloidin staining for contrast background of the hAECs monolayer. Finally, the constructs were stored at 4°C until imaged using a Nikon A1R fluorescence confocal microscope. Three ROI images (250μm x 250μm) at zone 1 (CCA) and zone 2 (Bifurcation to ICA and ECA) were captured from each construct using a Nikon A1R fluorescence confocal microscope.

### Statistical analysis

In each study, a minimum of three experimental replicates (n ≥ 3) were included, unless specifically noted otherwise. The results are presented as mean values accompanied by the corresponding standard deviation. Data from the experimental characterization underwent analysis using an unpaired t-test and one-way or two-way ANOVA statistical approaches (GraphPad Prism 9 software, United States). Tukey’s test for multiple comparisons was employed to further analyze distinctions between various groups. The significance threshold was set at a confidence interval of 95%, meaning that a p-value of less than 0.05 was considered statistically significant. For precise interpretation, the levels of significance were indicated as follows: *p < 0.05, **p < 0.01, ***p < 0.001, and ****p < 0.0001.

## Acknowledgments

The research is supported by the Medical Research Future Fund (MRFF 2016165, 2023977) awarded to ZYL; the Australian Research Council (FT180100157, DP200101658, DP230100721, IC190100026) awarded to YCT; and JAC is supported by the QUT Centre for Biomedical Technologies Scholarship and ARC DP200103942. This work was enabled using the Central Analytical Research Facility (CARF) at the Queensland University of Technology (QUT).

## Supporting Information

Supporting Information is available from the Wiley Online Library or from the author.

Received: ((will be filled in by the editorial staff))

Revised: ((will be filled in by the editorial staff))

Published online: ((will be filled in by the editorial staff))

---

This study introduces the first miniaturized, patient-specific carotid artery model created via 3D printing using GelMA with embedded vascular cells. Combining CFD, PIV, and flow perfusion, the model replicates anatomical-dependent hemodynamics and cellular responses. It accurately reflects shear stress effects and inflammation-prone zones, offering a valuable tool to study atherosclerosis and vascular remodeling.

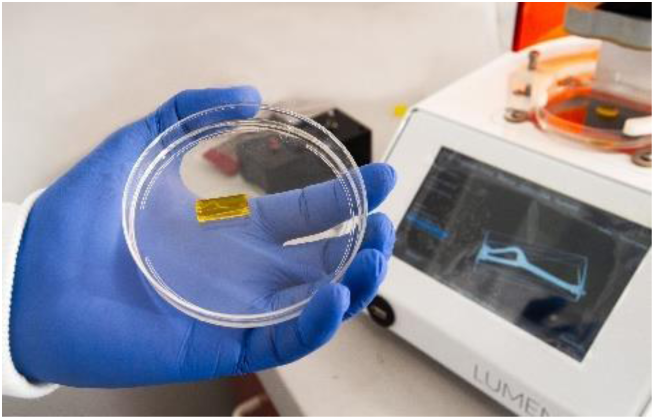

## Supporting Information

**Figure S1.**
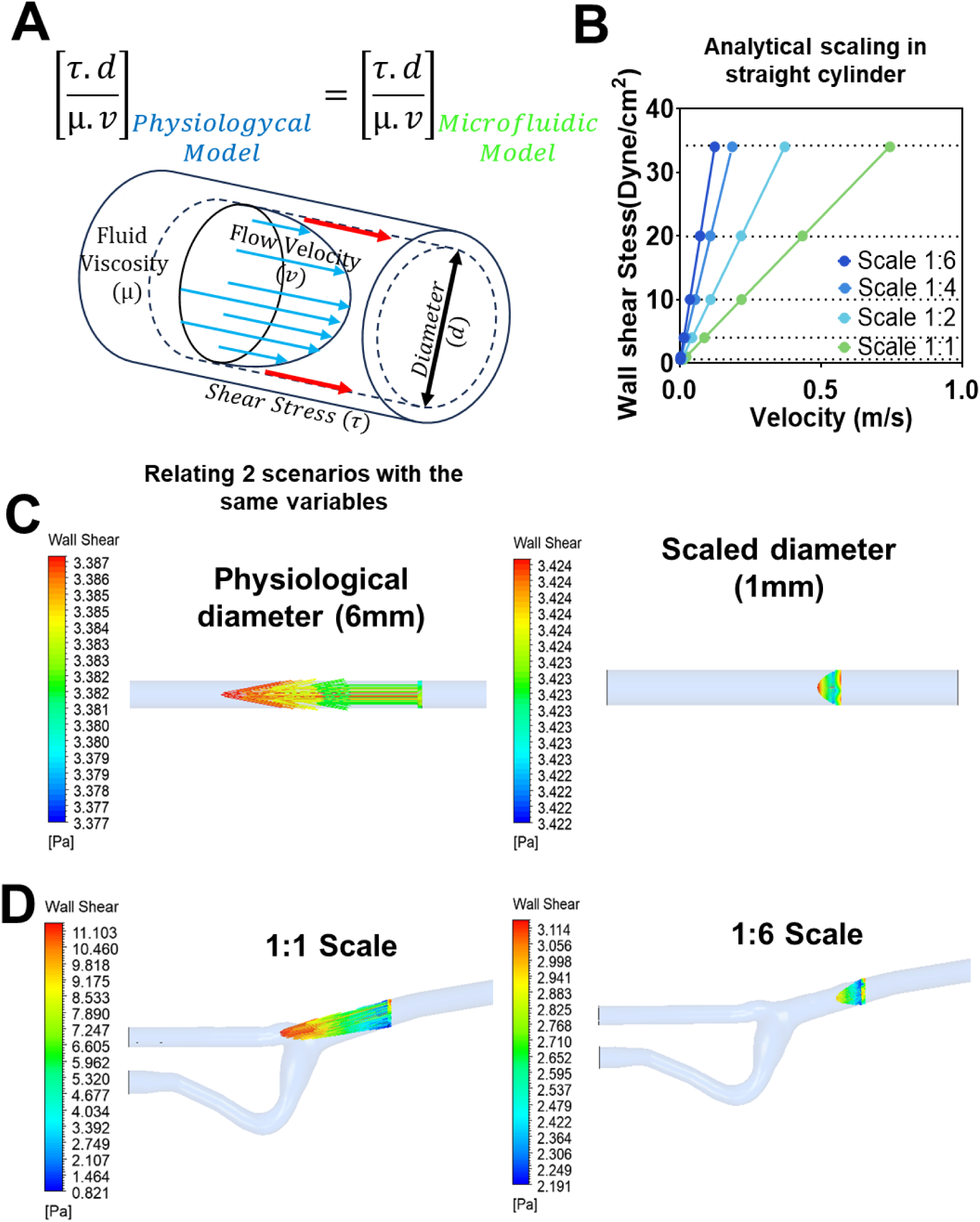
Dimensional analysis for scaling physiological flow velocity to a microfluidic model while maintaining constant shear stress. (A) Analytical similitude equation and schematic diagram illustrating the relationships between variables relevant to cylindrical fluid dynamics. (B) Equivalent wall shear stress (WSS) as a function of flow velocities in straight cylinders of 1:1, 1:2, 1:4, and 1:6 scales determined by analytical similitude analysis. (C) CFD simulations using analytically calculated velocities to compare wall shear stress distributions at the physiological scale (1:1) and the microfluidic scale (1:6) on simplified cylindrical models. (D) CFD simulations applied to the reconstructed carotid geometry, comparing wall shear stress at both scales.

**Figure S2.**
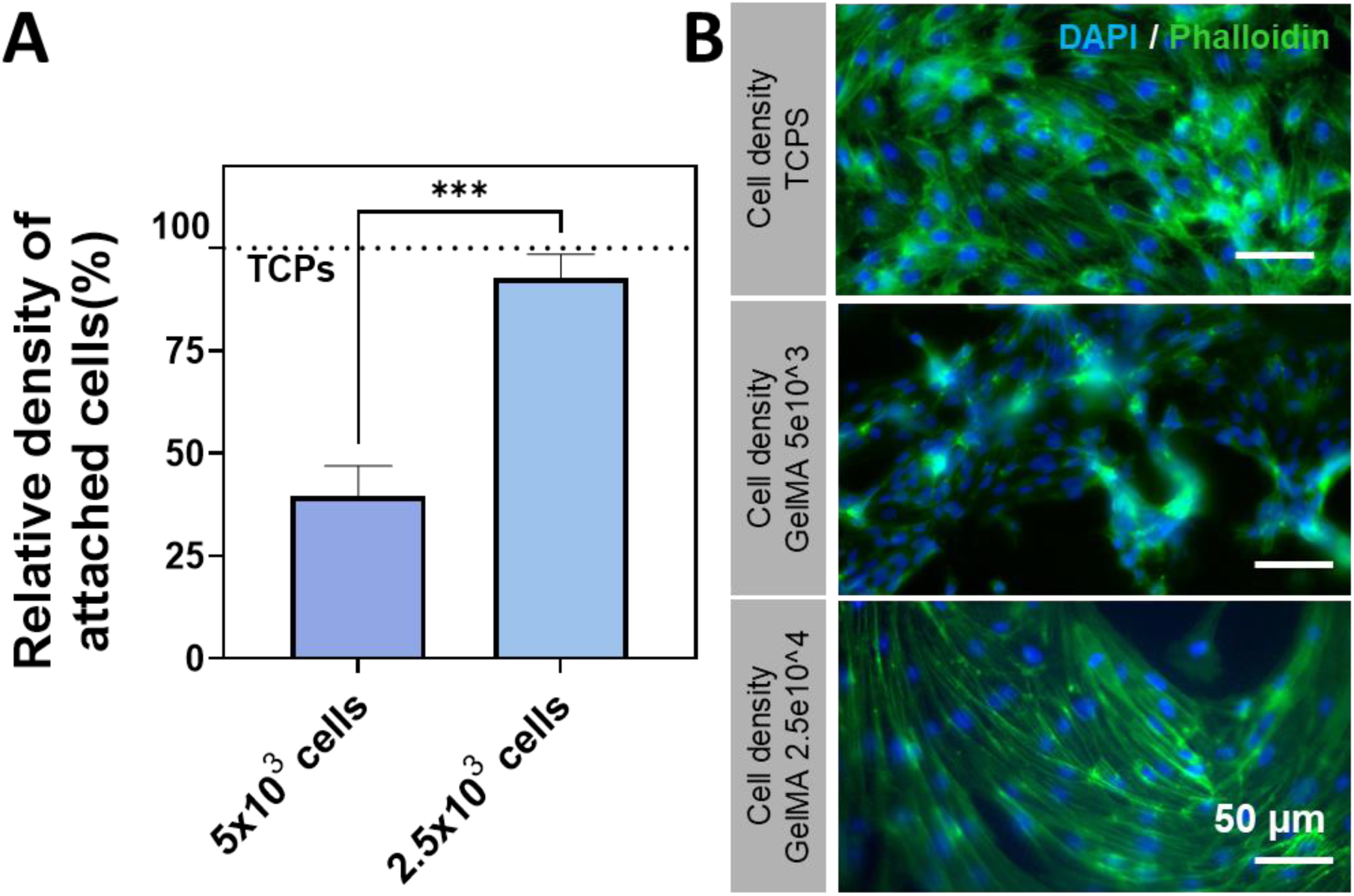
Effect of seeding density on HAEC attachment and cytoskeletal organization on GelMA. (A) Relative cell attachment density of GelMA substrates at standard TCPS seeding density (5,000 cells cm^-^²) and 25,000 cells cm^-^². (B) Fluorescence images of HAECs post-seeding, showing nuclei (DAPI) and cytoskeleton (Phalloidin) at densities of 5,000 and 25,000 cells cm^-^². Images were captured at 20× magnification, with a 50 µm scale bar. Statistical analysis was performed using a t-test. Data are presented as mean ± SD, n = 3. Statistical significance is denoted as ***p < 0.001.

**Figure S3.**
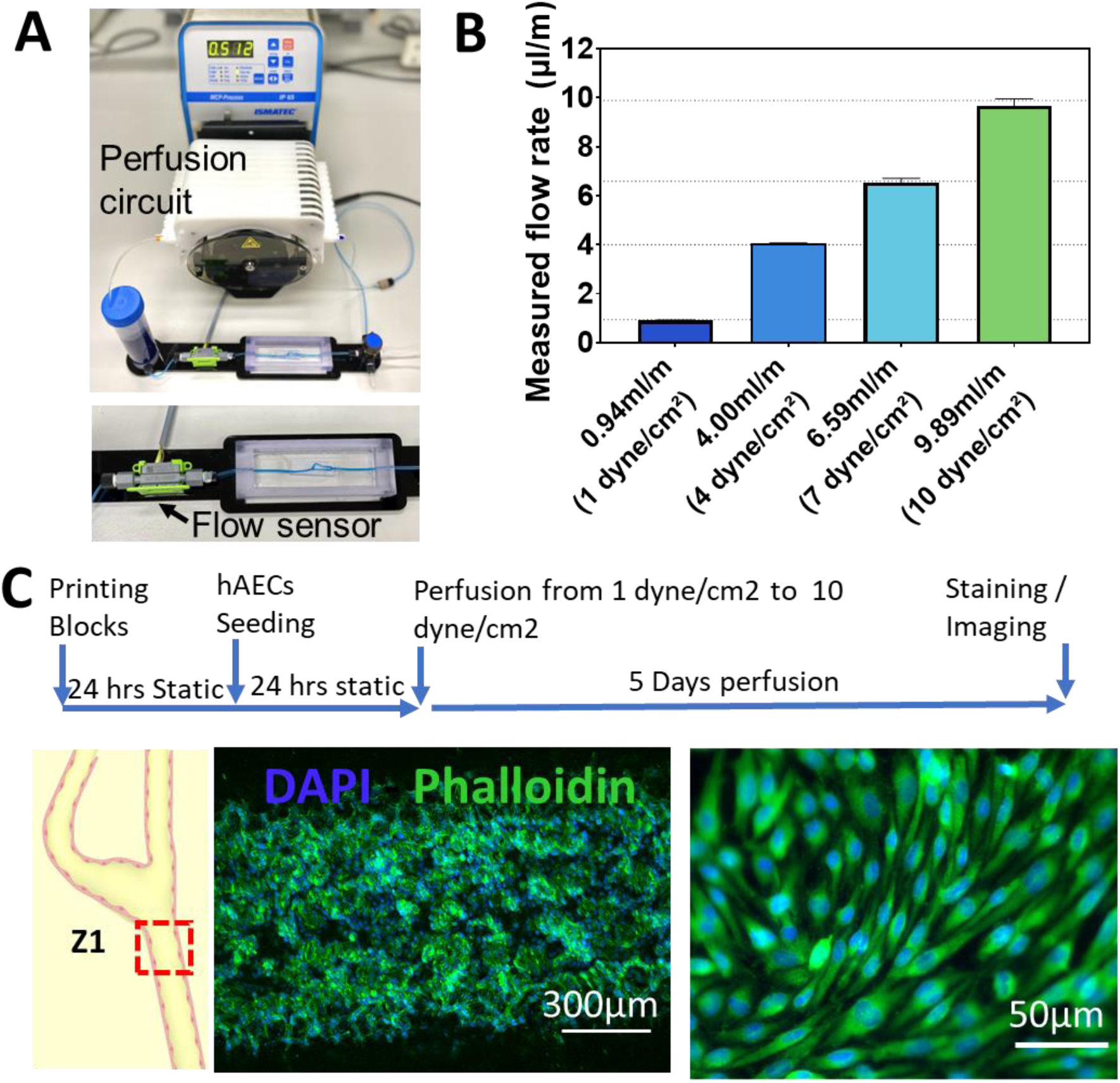
Evaluation of flow rate and hAEC attachment in GelMA-Collagen hydrogel channels under perfusion. (A) Experimental setup with an in-line flow rate sensor integrated into the perfusion circuit. (B) Real-time flow rate measurements corresponding to wall shear stresses of 1, 4, 7, and 10 dynes cm^-^². (C) Chronological progression of hAEC attachment during perfusion experiments, scaling shear stress from 1 to 10 dynes cm^-^². Fluorescence images of HAECs captured at 4× and 20× magnification, showing nuclei (DAPI) and cytoskeleton (Phalloidin) after 5 days of perfusion culture.

**Figure S4.**
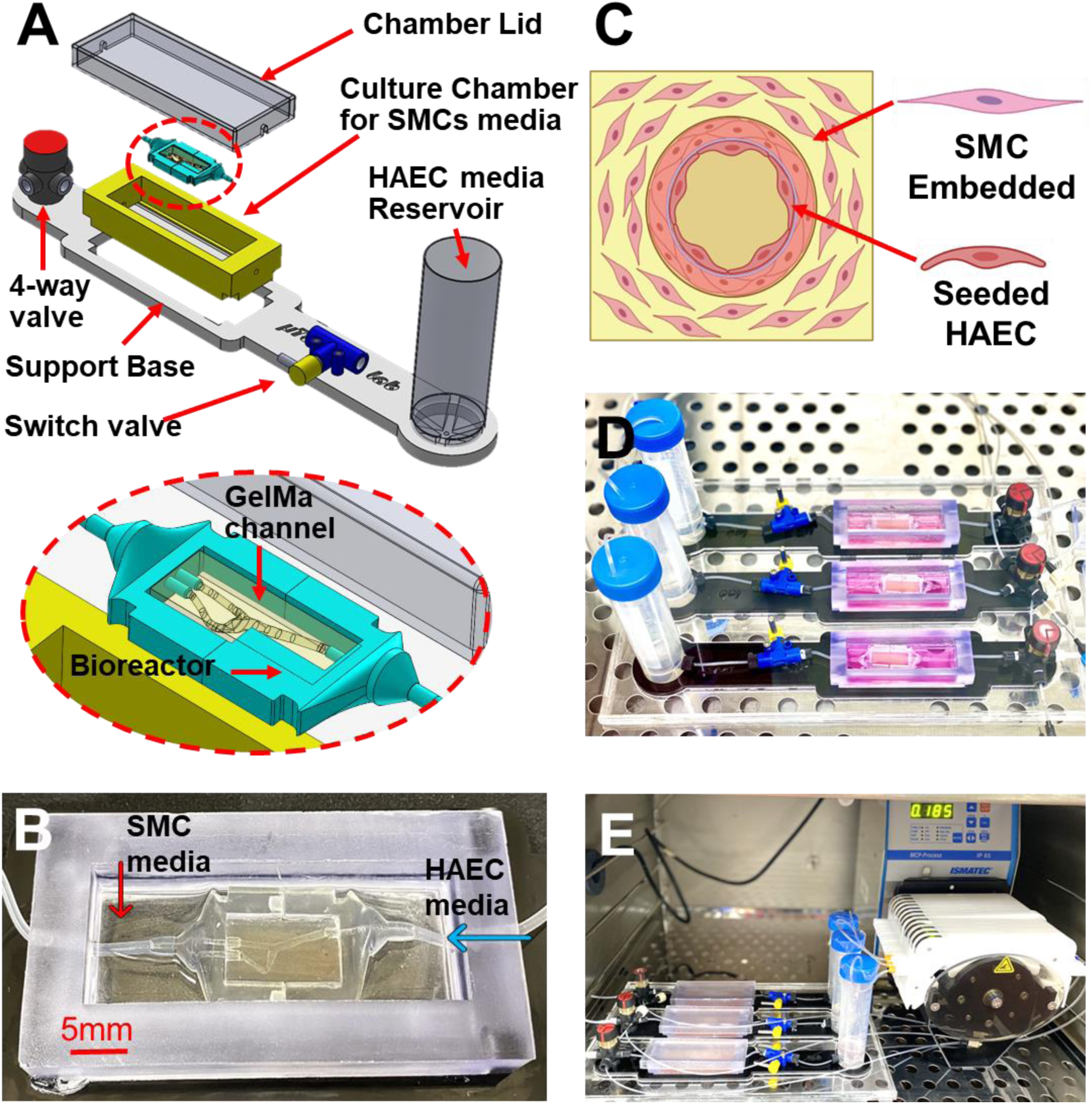
Perfusion system design for co-culture in 3D-printed GelMA-Collagen channels. (A) "World-to-chip" integration schematic and perfusion loop design, highlighting key components: bioreactor, enclosure chamber, base support, valves, and reservoir. (B) Bioreactor and enclosure chamber assembly, showcasing the dual media source configuration for feeding hAECs and hASMCs independently. (C) Cellular arrangement within the 3D DLP-printed hydrogel channel, with hASMCs embedded and hAECs seeded on the luminal surface. (D) Diagram of the co-culture perfusion loop under static conditions. (E) Perfusion setup illustrating dynamic conditions for the vascular construct.

**Figure S5.**
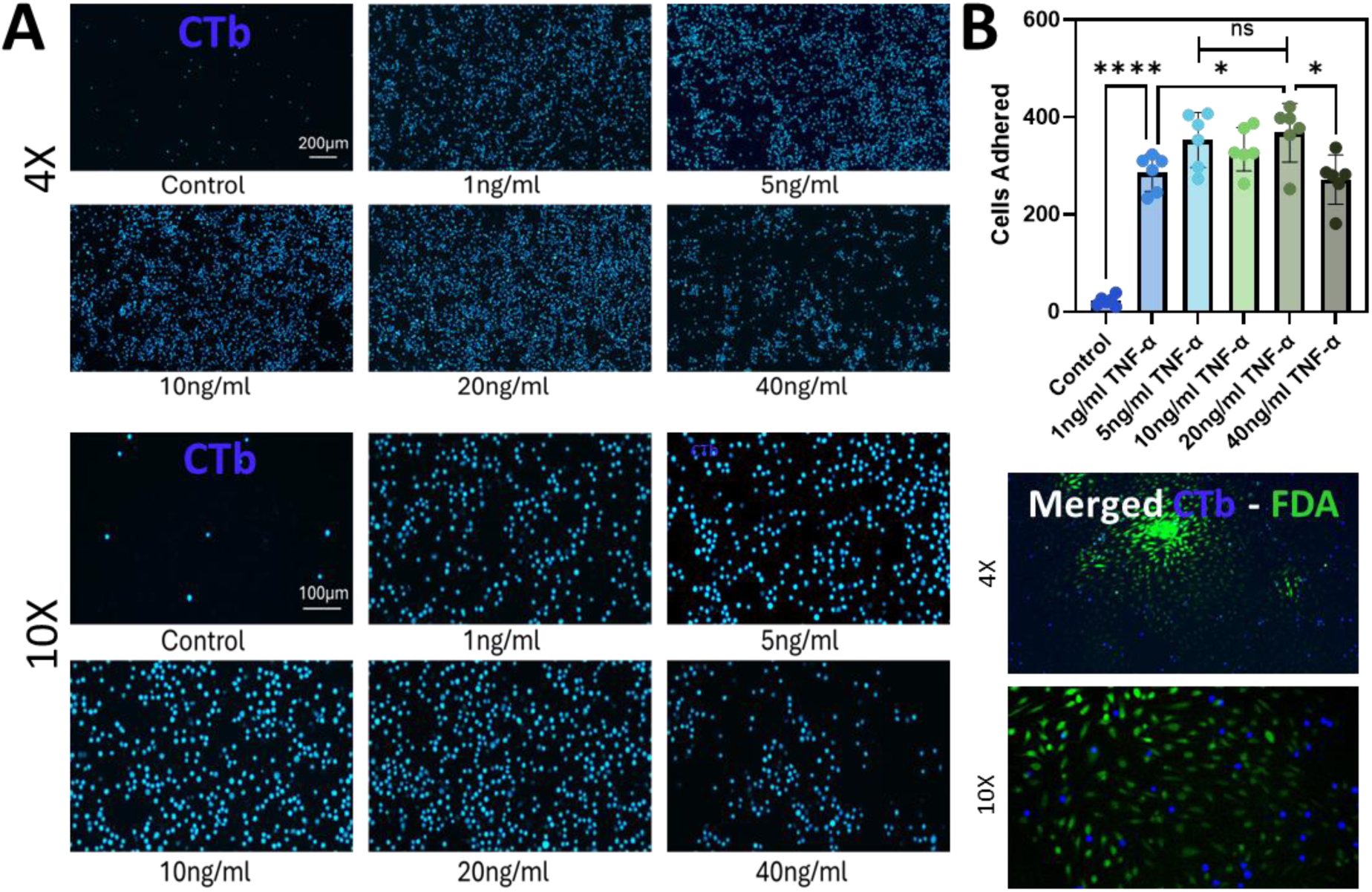
TNF-α concentration characterization for monocyte adhesion on 2D cultures. (A) Fluorescence imaging of adhered U937 monocytes (CTb-stained, blue) under varying TNF-α concentrations (0, 1, 5, 10, 20, and 40 ng mL^-2^). Images captured at 10× magnification (scale bar: 200 µm) and 4× magnification (scale bar: 100 µm). (B) Quantitative analysis of monocyte adhesion under different TNF-α treatments. Data presented as mean ± SD (n=3). Statistical significance determined using one-way ANOVA, with significance levels denoted as ****p < 0.0001; *p < 0.05.

